# SlicerMorph: An open and extensible platform to retrieve, visualize and analyze 3D morphology

**DOI:** 10.1101/2020.11.09.374926

**Authors:** Sara Rolfe, Steve Pieper, Arthur Porto, Kelly Diamond, Julie Winchester, Shan Shan, Henry Kirveslahti, Doug Boyer, Adam Summers, A. Murat Maga

## Abstract

Large scale digitization projects such as *#ScanAllFishes* and *oVert* are generating high-resolution microCT scans of vertebrates by the thousands. Data from these projects are shared with the community using aggregate 3D specimen repositories like MorphoSource through various open licenses. MorphoSource currently hosts tens of thousands of 3D scans of eukaryotes. Along with the data from similarly scoped projects such as 10kPhenomes, DigiMorph and many others, soon hundreds of thousands of specimens that represent biodiversity of extinct and extant organisms will be conveniently available to researchers. We anticipate an explosion of quantitative research in organismal biology with the convergence of available data and the methodologies to analyze them.

Though the data are available, the road from a series of images to analysis is fraught with challenges for most biologists. It involves tedious tasks of data format conversions, preserving spatial scale of the data accurately, 3D visualization and segmentations, acquiring measurements and annotations. When scientists use commercial software with proprietary formats, a roadblock for data exchange, collaboration, and reproducibility is erected that hurts the efforts of the scientific community to broaden participation in research. Another relevant concern is that ultimate derivative data from individual research projects (e.g., 3D models of segmentation) are shared in formats that do not preserve the correct spatial scale of the data.

In this paper, we present our effort to tackle challenges biologists face when conducting 3D specimen-based research. We developed SlicerMorph as an extension of 3D Slicer, a biomedical visualization and analysis ecosystem with extensive visualization and segmentation capabilities built on proven python-scriptable open-source libraries such as Visualization Toolkit and Insight Toolkit. In addition to the core functionalities of Slicer, SlicerMorph provides users with modules to conveniently retrieve open-access 3D models or import users own 3D volumes, to annotate 3D curve and patch-based landmarks, generate canonical templates, conduct geometric morphometric analyses of 3D organismal form using both landmark-driven and landmark-free approaches, and create 3D animations from their results. We highlight how these individual modules can be tied together to establish complete workflow(s) from image sequence to morphospace. Our software development efforts were supplemented with short courses and workshops that cover the fundamentals of 3D imaging and morphometric analyses as it applies to study of organismal form and shape in evolutionary biology, and extensive links to the existing tutorials are provided as supplemental material.

Our goal is to establish a community of organismal biologists centered around Slicer and SlicerMorph to facilitate easy exchange of data and results and collaborations using 3D specimens. Our proposition to our colleagues is that using a common open platform supported by a large user and developer community ensures the longevity and sustainability of the tools beyond the initial development effort.

## INTRODUCTION

A keyword search on the US National Library of Medicine Pubmed database for keywords (**ANATOMY** OR **MORPHOLOGY**) AND **microCT** results in over 6000 indexed publications for non-human animal species. Given the emergence of other 3D imaging modalities such as stereophotogrammetry (3D surface reconstruction), MRI, 3D confocal and light-sheet microscopy, the real tally of papers using some sort of 3D imaging modality are in the tens of thousands, and most of these papers are published in the last five years. This shows the impact of 3D imaging methods on the analysis of anatomical structure whether the question is developmental, evolutionary or genetic in nature. Going forward, we expect this trend to continue even more strongly thanks to the convergence of availability of large numbers of datasets and new methods to analyze them.

Large scale, publicly funded biodiversity digitization efforts such as *ScanAllFishes* and *openVertebrates* [1,2] and others are documenting the 3D anatomy in unprecedented detail and in numbers. Additionally, aggregate specimen repositories like MorphoSource facilitate finding datasets across projects rather conveniently [3]. Coupled with more biomedically oriented 3D imaging archives, such as FaceBase [4], there is now an enormous and expanding amount of 3D data on biodiversity, particularly of skeletal structures. Thanks to the amenability of their mineralized skeletal structures to the X-ray based imaging methods, currently there is a clear vertebrate bias in the existing data. However, with the ever-increasing resolution of imaging systems and improvements to contrast enhancement both by digital [5] and traditional (e.g., radiopaque contrast agents) methods [6–9], it is only a matter of time before other non-vertebrate multi-cellular biological systems from all scales of life are represented in 3D specimen repositories.

In addition to the increasing availability of 3D biodiversity data, we are also experiencing an increase in the availability of methods to analyze them. Traditionally, quantitative inquiries into organismal shape and size of biological systems relied on morphometric methods based on linear measurements acquired directly from specimens. In the last couple decades, this traditional approach has been supplemented with geometric approaches, in which the input into the analysis is the Euclidean coordinates of ‘landmarks’ instead of distances between them [10]. Downstream analyses preserve this geometric arrangement of landmarks while adjusting for uniform size differences, and minimizing the difference between forms. These ‘Geometric Morphometric Methods’ (GMM) are now applied in many subdomains of biological sciences that use phenotypic variability in organismal form to answer questions, but particularly in evolutionary and developmental biology, as well as quantitative and population genetics fields. There is a rich literature on both theoretical and practical applications [11–17]. GMM relies on the expert annotation of biologically homologous landmarks by the investigator. Depending on the anatomical system being studied, as well as the type of the investigation, this requirement can be relaxed by using geometrically, instead of biologically, homologous landmarks to model and represent the underlying complex topology [18]. These geometrically constructed landmarks can be called semi-landmarks, or pseudo-landmarks depending on how they are generated. In addition to the explicitly user-generated landmark-driven geometric approaches, there are now methods that accomplish the correspondence of these landmarks across samples with or without user guidance [19–23]. The deep-learning class of machine learning methods also offers the potential to automatically detect and place landmarks, provided sufficiently large training and validation datasets exist[24,25].

In sum, quantitative analysis of organismal form is going through an “interesting time”. It is interesting because while the wealth of data and analytical methods offers promise of exciting breakthroughs, working with these datasets are typically fraught with challenges for most biologists. It involves tedious tasks of format conversions across a myriad of proprietary and open file formats, preserving spatial scale of the data accurately, 3D visualization and segmentation of structure of interest, acquiring measurements and annotations, and finally sharing the derivative datasets and results with the community. These steps may involve serial use of multiple software tools to accomplish the common steps of image formation, enhancement, visualization and analysis in most 3D image processing workflows (Figure 1).

**Figure 1.**
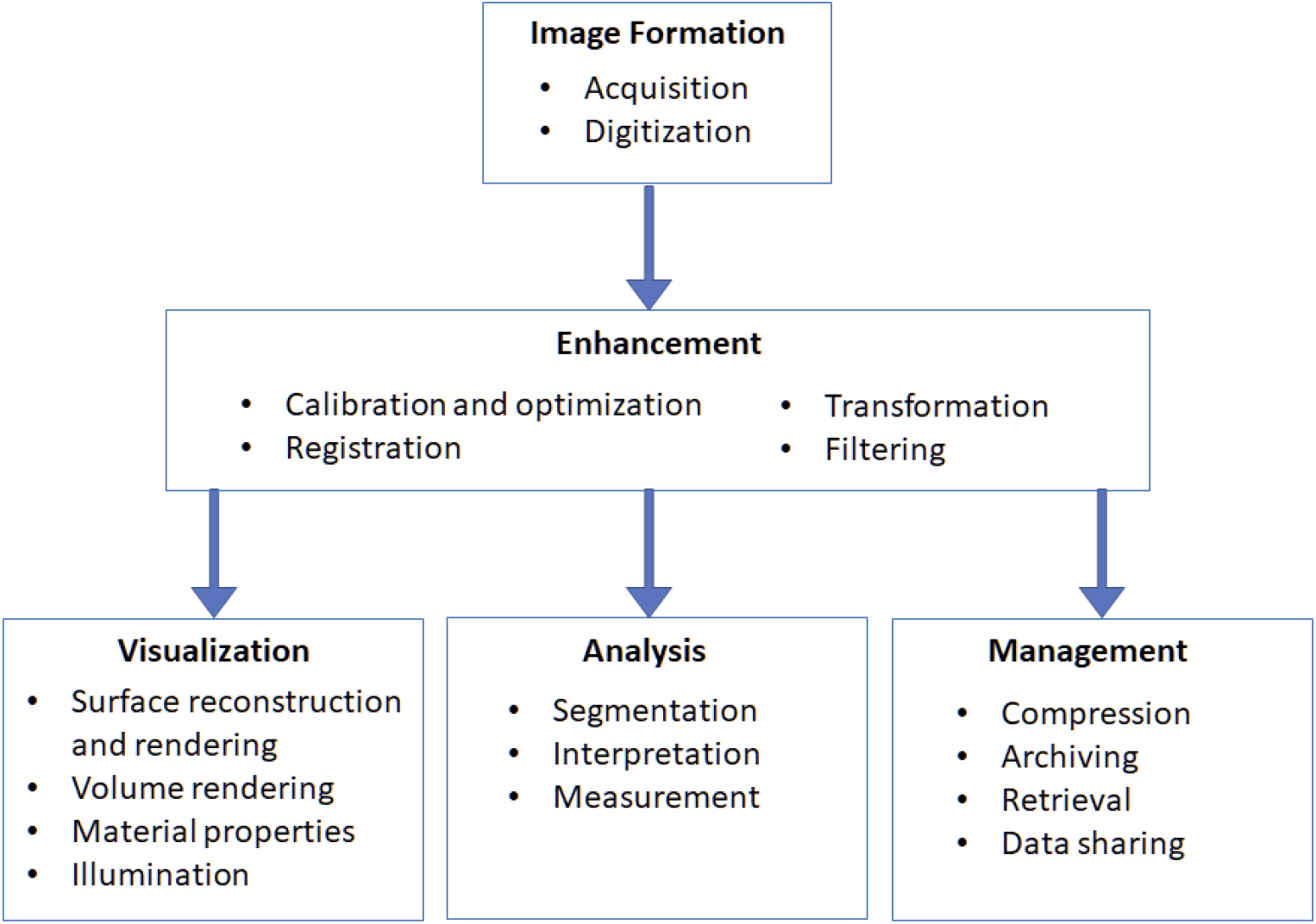
Steps in a 3D imaging workflow.

3D Slicer (Slicer) is an open-source biomedical visualization and image analysis software developed in the last 20 years predominantly by members of neuroimaging and surgical planning communities that share many of the concerns and frustrations currently felt by the biologists working with 3D specimen data [26–28]. Having a free, but feature-rich, open and extendable software to visualize data across projects consisting multiple investigator teams and a mix of operating systems (Windows, MacOS and Linux) was and remains a common motivating problem. Through the years, Slicer has grown into a mature ecosystem that can handle all the tasks associated with 3D image analysis (Figure 1), except for scanning. Core functionality of Slicer offers a complete solution for 3D visualization (both volume and surface based rendering), linear and nonlinear spatial transforms, manual and semi-automatic segmentation tools, 3D landmark (fiducial) and other measurement digitization, numerous image processing and enhancement filters from SimpleITK that are specifically 3D in nature, data type conversion (e.g., from 3D models to segmentations), plotting and tabular data representation, and a built-in Python3 environment that includes common libraries such as numpy and scipy. Functionality that is not available in the core application can be developed through an extension mechanism. Additionally, most (but, due to binary dependencies, not necessarily all) Python3 libraries from the PyPi repository can be installed into the integrated Python environment using the standard Python pip utility. As of writing, the combined downloads of the previous and current stable versions (v4.10 and v4.11 respectively) since November 2018 exceed 270,000 worldwide [29]. There are over 10,000 publications indexed in Google scholar that cite 3D Slicer. Slicer has also been adapted to a wide range of use case, such as the substantial SlicerAstro effort [30,31]. Thus, Slicer has a vibrant ecosystem, thanks in particular to its extensible code base and its reliance on proven, open-source libraries such as Visualization Toolkit (VTK), and Insight Toolkit (ITK), an active global developer community, and effective user support through a forum that is followed actively over 3,500 subscribers and averages 250 posts weekly [32].

The Slicer community maintains the “app store” for Slicer extensions, currently with well over 100 extensions. These extensions provide domain-specific functionality so that users can customize Slicer to their needs without introducing special case code in the core of Slicer that would complicate it for other use cases. For example, the SlicerDMRI extension provides extensive custom code to support diffusion MRI data processing and tractography that is only useful for users working in that field. Slicer extensions can be written in C++ or Python, and are built each night from source to be compatible with the corresponding pre-release Slicer preview builds for the supported OS platforms. Extensions are also built nightly for the current Slicer release, allowing extension developers to develop new functionality against a stable platform if they don’t need features that are only available in the current nightly preview. Some Slicer extensions include SlicerIGT for Image-Guided Therapy [33], SlicerCMF for craniomaxillofacial surgery [34], and SlicerRT for radiotherapy research [35].

## SLICERMORPH

SlicerMorph is an extension built onto the current stable version of Slicer (r29402) and will not work with earlier stable versions (e.g., v4.10.2, r 28257) as these lack necessary features that SlicerMorph relies on. Official method of installing SlicerMorph is using the built-in extension manager in Slicer (See SOM #1). On all three major operating systems, Slicer installs by default into the user space, and does not require elevated (admin) user rights on the computer. MacOS and Windows users looking for portability (e.g., running from a thumb drive) can opt to use a prepackaged version we maintain independently at http://download.SlicerMorph.org. This version comes with all extensions and python libraries preinstalled and requires only expanding the archive to a user accessible folder. Source code of all SlicerMorph modules can be obtained at https://github.com/SlicerMorph/SlicerMorph.

After the install, SlicerMorph extension registers a new section called SlicerMorph preferences in the Application Settings of Slicer. SlicerMorph settings are off by default, but can be enabled by the user. Tab contains the SlicerMorph specific customization of Slicer’s settings. These include overriding some of Slicer’s default settings such as increasing unit precision, making orthographic rendering the default choice, enabling rulers in slice and 3D views, as well as unique customizations such as disabling compression (for faster read and write operations of large datasets), setting the default model output format to PLY, and registering custom keystrokes for faster landmarking and segmentation operations. These settings are stored in an editable python script file, in which users can add into their specific shortcuts and actions using the examples provided. Installing SlicerMorph also registers new sample datasets with the Sample Data module. Users new to the Slicer platform who need more instruction on how to use Slicer and extension mechanism should refer to SOM #2, which also provides a link to the official SlicerMorph documentation. An abbreviated glossary of technical terms used in this paper and in Slicer platform is provided as SOM #3.

SlicerMorph also imports a number of other Slicer extensions which provide complementary functionality such as importing 3D volumes from other software (RawImageGuess), additional segmentation filters (SegmentEditorExtraEffects), 3D shading control (Lights), creating rigid and warping transformation from two landmark sets that can be applied to any data node (FiducialRegistration) and others.

The modules within SlicerMorph extension can be grouped into three main categories:

1. Input/Output
2. Geometric Morphometrics
3. Utilities

Below, we provide a brief description of what each SlicerMorph module does. Later, in the workflow section, we explain the reasons they were developed and how they differ from other modules in Slicer that offer similar functionality.

### Input/Output modules

1. **ImageStacks:** A general purpose tool to import non-DICOM image sequences (png/jpg/bmp/tiff) into Slicer. Users can specify voxel size, select partial range, downsample (50%) along all three axes, load every Nth slice (skip a slice), or reverse stack order (to deal with left/right mirroring of the specimen due unknown slice ordering convention of the data). ImageStacks also reports the estimated memory requirement to load the full and downsampled data set based on the image data type, so that users can be informed about the memory requirements.
2. **SkyscanReconImport:** Imports an image stack from the widely used Bruker/Skyscan microCT reconstruction software (Nrecon). The only input to the module is the reconstruction log file (*_Rec.log) that was generated by the Nrecon software. Necessary information about correct voxel spacing, volume filename prefix, dimensions, are read from the log file, and if there is a discrepancy between image data and log file entries, an error is generated. Left/right mirroring of the specimen is avoided, as the correct image ordering for Bruker/Skyscan is built into the module.
3. **MorphoSourceImport:** Provides a direct way to query and retrieve data from the aggregate specimen repository MorphoSource into the current Slicer scene. Returned results are restricted to the 3D models that have been released under an open-access license. Users need to register and acquire a username from the MorphoSource website.
4. **ExportAs:** This plugin to the existing Data module of Slicer provides a one-click (right-mouse context menu) export of any data node loaded into the Slicer scene at the time. While the regular Save dialog box is suggested for complex workflows involving multiple different data types, ExportAs is convenient for users with simpler workflows that involve few data nodes (e.g., a 3D model and accompanying landmark set).

### Geometric Morphometrics modules

1. **Generalized Procrustes Analysis (GPA):** Performs GPA with or without scaling shape configurations to unit size, conducts principal component analysis (PCA) of GPA aligned shape coordinates, provides graphical output of GPA results and real-time 3D visualization of PC warps either using the landmarks of mean shape, or using a reference model that is transformed into the mean shape. Visualization of 3D shape deformation of the reference model can be exported as a video clip. The input into the module is a folder path containing a number of landmark files stored in Slicer’s FCSV format and, optionally, a 3D model and accompanying set of landmarks to be used as reference model in 3D visualization of PCA results.
2. **CreateSemiLMPatches**: Provides triangular patches of semi-landmarks that are constrained by three fixed anatomical landmarks. The input into the module is a 3D model and its accompanying set of fixed landmarks, and users generate and visualize the patches by specifying triplets of fixed landmarks that form a triangle.
3. **PseudoLMGenerator:** This module uses the 3D model’s geometry (or alternatively a spherical surface) to create a dense-template of pseudo-landmarks. The landmark placement is constrained to the external surface of the mesh. Points sampled on the template are projected to the mesh surface along the surface normals of the template. Projected points are then filtered to remove those within the sample spacing distance, improving regularity of sampling. This module can be used to generate a large number of points on a 3D model that can serve as a landmark template for additional samples in the dataset.
4. **Automated landmarking through point cloud alignment and correspondence analysis (ALPACA)**: ALPACA provides fast landmark transfer from a reference 3D model and its associated landmark set to target 3D model(s) through point cloud alignment and deformable registration. The optimal set of parameters that gives the best correspondence between a single pair of reference and target models can be investigated in the single alignment mode. Once an optimal set of parameters are found, these can be applied to a number of 3D models in batch mode.
5. **Auto3dgm**: Auto3dgm is a Python3 implementation of a previously published method for comparative analysis of 3D digital models representing biological surfaces [20,36]. Unlike other three-dimensional GMM, auto3dgm uses an automated procedure for placing landmarks on surfaces that might be devoid of anatomical landmarks, or too dissimilar to use template-based methods. The input to the software is a folder containing 3D models to be placed pseudo-landmarks on. Users then have the option to specify the number of pseudo-landmarks requested, and number of iterations to be run. The results of 3D models alignment, and resultant pseudo-landmarks can be visualized, and if satisfactory, can be analyzed with the GPA module described above. Auto3dgm can also be acquired independently of SlicerMorph as a separate extension. Because it is maintained outside the SlicerMorph project, source code for auto3Dgm is available from https://github.com/ToothAndClaw/SlicerAuto3dgm. Note that auto3Dgm uses Mosek, a highly optimized, proprietary mathematical solver, which requires a license to run. A free academic license for Mosek can be acquired from the vendor online.
6. **MarkupEditor**: A module that enables selecting and editing subsets of landmarks on a model by drawing an arbitrary closed curve in the 3D viewer using right-click context menu. Landmarks must be visible from the current camera view angle. Selected landmarks can be removed from the active fiducial list or copied into a new one. This module is useful to quickly identify and remove unwanted points in pseudo- or semilandmark sets, or group landmarks into classes for further analyses (e.g., for modularity and integration).

### Utility Modules

1. **Animator**:A basic keyframe-based animator of 3D volumes. Supports interpolation of regions of interests, rotations, and transfer functions for volume rendering. Output can be either as a sequence of frames or a movie in mp4 format.
2. **ImportFromURL:** Imports any data with a public URL, as long as it is in a format supported by Slicer.
3. **ImportSurfaceToSegment**: Imports any 3D model format read by Slicer as a segmentation node and prompts the user to edit it using the Segment Editor module.
4. **MorphologikaLMConverter**: Imports a legacy Morphologika formatted landmark file that may contain multiple subjects and exports a landmark file for each individual to a directory specified by the user in Slicer’s. FCSV format.
5. **IDAVLMConverter**: Imports the legacy IDAV landmark editor files into the existing Slicer scene. One data file per specimen is expected (pts format).
6. **PlaceSemiLMPatches:** A utility module that applies the generated connectivity table from the CreateSemiLMPatches module to other 3D models with the same set of anatomical landmarks. A new set of semiLandmarks are sampled directly from the new models using the specified connectivity map. Input are: the connectivity table from CreateSemiLMPatches, a directory with the list of 3D models that semi-landmarks will be placed on, and associated anatomical landmark files (in FCSV format). Models and their corresponding FCSV files should have the same filename prefix.
7. **ProjectSemiLM:** A utility module to transfer a template of semilandmarks to new 3D models using Thin Plate Splines (TPS) warp. Requires a set of corresponding anatomical landmarks in the template and target models. Required inputs are: a base 3D model with its anatomical landmark and the semi-landmark template sets used as reference; a directory with the list of 3D models that semi-landmarks will be projected on and their associated anatomical landmark files (in FCSV format). Model and the corresponding FCSV files should have the same filename prefix.
8. **VolumeToModel:** A convenience module to segment a 3D volume by specifying a threshold for intensity and then export the segmentation as a 3D model into the current Slicer scene. Useful for directly extracting models from scans that contain only dry skeletons.
9. **SegmentEndoCranium:** Automatically segments the endocranial space in the 3D volume of a vertebrate skull using a combination of effects from the Segment Editor module. User needs to specify the diameter of the largest hole that needs to be plugged during the endocast creation.

## WORKFLOWS IN SLICERMORPH

SlicerMorph offers a complete workflow from raw data import to morphometric analysis. We split the workflow into four different tasks: Data Import, 3D Visualization, Segmentation, GM Data Collection and Analysis. Because Slicer is extensible through both extensions and by importing new Python3 libraries, there is usually more than one way of accomplishing the same task. Workflows presented here are by no means the only way to process data in Slicer, but that are robust methods based on the feedback we received from over a hundred participants in our workshops over the past three years. The workflows assume the user installed SlicerMorph as described above.

## 1. DATA IMPORT WORKFLOW

3D scans of biological specimens exist either as 3D models or 3D volumes. Note that none of the common file formats were created with the use case of biological specimens in mind, so they do not natively record essential metadata such as spatial dimension units, specimen provenance, species, or acquisition methods, so care must be taken to record such data when using these commodity formats. Below we discuss specifics of importing each type of data, and their potential pitfalls, particularly for quantitative morphology studies. Figure 2 provides the summary of the suggested workflow.

**Figure 2.**
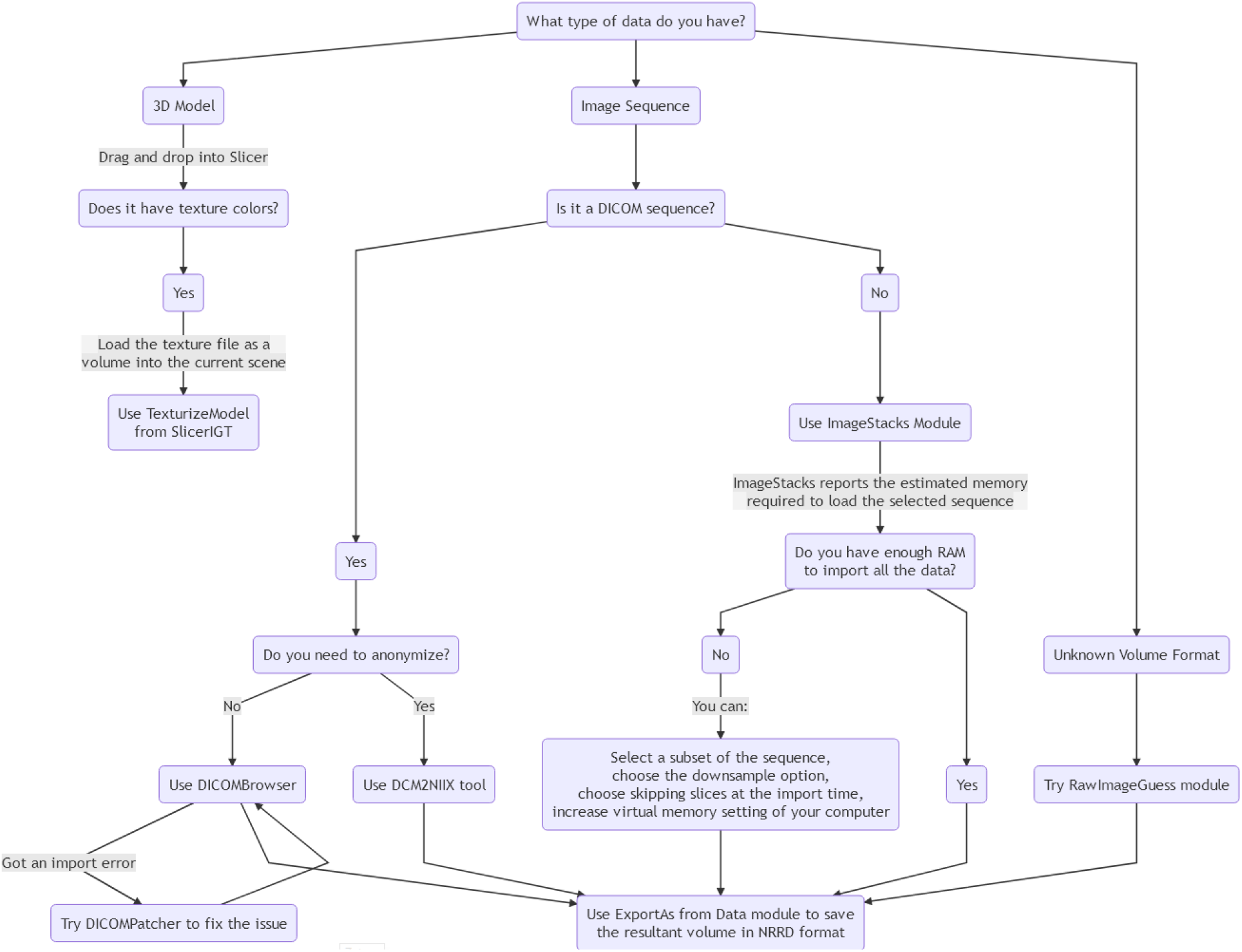
SlicerMorph data import workflow.

### 3D Models

Slicer natively supports common 3D model formats OBJ, STL, PLY and VTK. These formats can be directly read into Slicer by dragging and dropping them into the application window or using the Open Data dialog box. If the models contain texture, the bitmap texture file can be loaded into the scene as a volume, and then the TexturizeModel module of the SlicerIGT extension can be used to apply the texture to the imported model.

### DICOM Image Sequences

Datasets from volumetric image acquisition, such as microCT or MR, are usually saved as an image sequence. One format, Digital Imaging and Communications in Medicine, or DICOM, is the international standard for medical images and related information. Slicer’s DICOM module provides the functionality to parse and import DICOM sequences. Since DICOM is used for a wide variety of purposes, the DICOM module supports a plugin architecture so that extensions can provide custom loaders for a specific data use case; for example, the SlicerRT extension provides plugins for a number of key radiotherapy data types such as dose maps and beam specification plans. DICOM is a very metadata rich format, and may contain extensive additional information about the subject, as may be the case if the data is clinical in nature (e.g., scans from veterinary clinics). Particular care should be paid to subject privacy when working with such datasets, and an anonymization step might be necessary. Slicer itself does not provide DICOM anonymization, but several open source tools are available such as DicomCleaner (PixelMed) and CTP (Radiological Society of North America). It is also possible that while the image sequence might nominally be in DICOM format, it may not actually be compliant with the standard, which Slicer closely adheres to. We observed this to be frequently the case when DICOMs are secondarily exported from other formats. In such cases, a patching step might be necessary to successfully import these datasets into Slicer. Patching can sometimes be accomplished by using the DICOMPatcher module of Slicer. Alternatively, external tools such as DCM2NIIX can be used for this step [37].

### Non-DICOM Image Sequences and their pitfalls

3D scans of specimens stored in non-DICOM image sequences are also common. These might be generated by research microCT, optical projection or coherence tomography scanners and 3D microscopes such as confocal, or light-sheet. When sequences of 2D bitmap formats like JPG, BMP, PNG or TIF are used to represent 3D volumes, information about the voxel spacing needs to be stored externally, since, unlike DICOM, these formats lack a standard representation of this data. Decoupling of this critical piece of information about the scale from the primary imaging data can potentially lead to loss of information, and should be avoided. 3D volumetric scans are better stored in formats that preserve the scale and coordinate system of the data. An open-source alternative is Nearly Raw Raster Data (NRRD), which is a library and file format designed to support scientific visualization and image processing involving N-dimensional raster data. Besides dimensional generality, nrrd is flexible with respect to type (8 integral types, 2 floating point types), encoding of written files (raw, ascii, hex, or gzip or bzip2 compression), and endianness. I/O libraries and tools for NRRD are available in common programming languages and other image analysis tools such as ITKSnap, FiJi, Python and R. NRRD is the default 3D volume format for Slicer.

The SlicerMorph module ImageStacks provides a one-step interface to import non-DICOM image sequences. It expects a user selection of files that are consecutively named. While Slicer’s default data load dialog box also accepts these formats, the resultant data node is a vector (RGB color) volume and requires conversion to scalar (single channel) volume for use with most of the rendering or segmentation features of Slicer. Additionally, ImageStacks provide options to specify original voxel spacing and to downsample the images at the time of import. It also reports the amount of memory necessary to import the full image sequence. At this point a user can decide whether their system has sufficient memory to handle the data they are trying to load and consider their options. Typical workflows, particularly segmentation, require physical memory that is several times the size of the input volume. Another drawback of non-DICOM image sequences is that there is no specific convention about how the ordinal values of slice numbers relate to the 3D volume composition; in other words it is arbitrary whether the lower slice numbers represent the top or the bottom of the stack. Incorrect ordering of the image sequence may cause a mirror reflection of the specimen. To deal with this issue, ImageStacks module has an option to reverse the ordering of the image sequences.

The Bruker Skyscan line of desktop microCTs are relatively common in biology labs. We have developed a specific import module, **SkyscanReconImport**, that reads the accompanying log file from the reconstruction software and correctly imports the image sequence with correct spacing and orientation. In cases where the image sequence comes unmodified from these scanners, users will benefit from using SkyScanReconImport instead of ImageStacks, however this module lacks the downsampling options offered by ImageStacks. SkyscanReconImport has been tested up to v1.7.0.4 of Bruker Nrecon software. Where sufficient information is available, additional vendor specific import modules can also be developed within SlicerMorph.

To some extent, it is possible to directly import volumetric data from proprietary formats of commercial biomedical visualization software such as VG Studio or Avizo into Slicer. The success rate is dependent on the amount of information available for the format. Prior knowledge of volume dimensions, data type, endianness (if applicable), header size and whether the image is compressed or not greatly increase the success rate. For uncompressed volumes, specifics of image data can be directly entered into (or interactively explored through) the user interface of the **RawImageGuess** module. If the compression format is known and it is one of the algorithms supported by NRRD, the user can attempt to generate a detached NRRD header that describes the data and use that to import the volume into Slicer. But when possible, exporting the image data into NRRD (or other Slicer compatible 3D volume format) from the original software is the least error-prone option.

Once the 3D volume is successfully imported, orientation and the correct scale of the data is verified, users should save their data immediately in NRRD format to avoid repeating data import and to preserve image scale and geometry.

## 2. 3D VISUALIZATION WORKFLOW

The only addition by SlicerMorph to the already powerful 3D visualization capability of Slicer is the Animator module, which will be introduced below. Any 3D model imported into the Slicer is immediately displayed in the 3D renderer window. Color and material properties, and model surface representation (point cloud, shaded with or without edges) are controlled through the DIsplay options of the Models module.

3D volumes in Slicer can be visualized using the Volume Rendering module (Figure 3). Volume rendering, also known as volume ray casting, is a visualization technique for displaying image volumes as 3D objects directly, without requiring segmentation and surface extraction. This is accomplished by specifying color and opacity for each voxel, based on its image intensity. For medical imaging modalities (i.e., CT and MR), Slicer provides several preset mapping parameter sets that highlight structures of interest (bone, soft tissue, air, fat). Users have the option to fine tune these or create a new mapping from scratch under the Volume Property setting. For advanced effects such as data-based cutouts, users have the option to provide custom GPU shader code using the SlicerPRISM extension [38]. Both CPU and GPU based ray casting methods are implemented in Slicer. A dedicated GPU greatly increases the rendering performance, particularly for multi-gigavoxel volumes. However, for the GPU based rendering to perform, the entire 3D volume needs to fit into the GPU memory, as Slicer does not automatically downsample the volume to match the hardware capability. Users can do this for themselves with the CropVolume module. Multiple 3D volumes can also be rendered simultaneously in the same 3D renderer window by choosing the MultiVolume Raycasting (still marked as an experimental feature as of the 4.11 version). Users can specify a region of interest (ROI) to render only a portion of the volume, or adjust its material properties. More fine-grained control over shading can be achieved through the Lights module.

**Figure 3.**
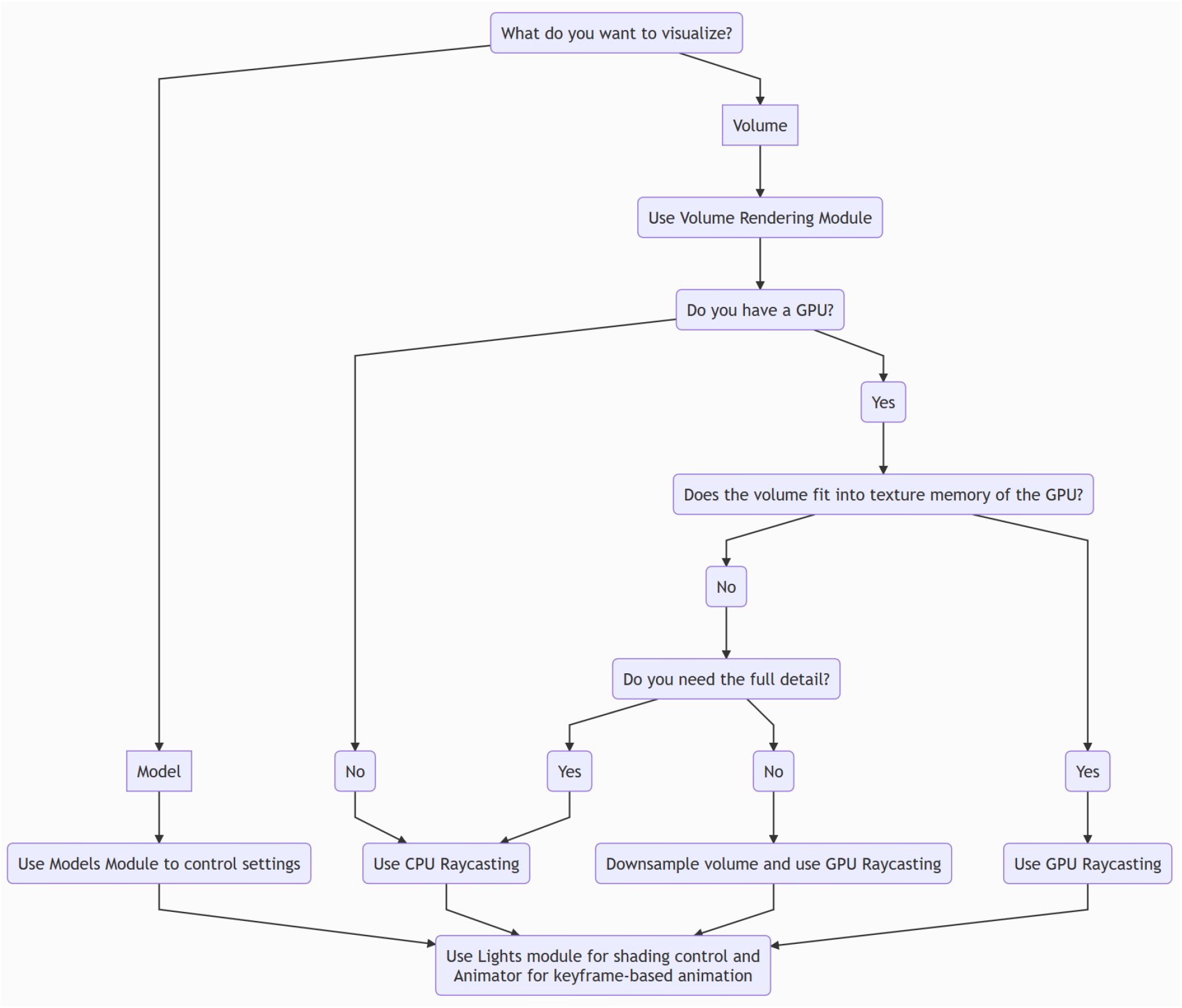
Data Visualization workflow in SlicerMorph.

### Animator

This SlicerMorph specific module provides simple keyframe-based animation capability in Slicer. Users can specify three types of keyframe actions: a starting and ending volume property (e.g., one that begins with highlighting the surface of the specimen, and ends only showing internal structures); starting and ending cropping ROIs (e.g., to virtually section the specimen from top to bottom); and camera rotation at a specified rate around a selected axis. The Animator module supports a plugin architecture so that Python programmers can add custom effects. The Animator module interpolates between keyframes and updates the rendering accordingly based on the timetrack of each action. While only a single ROI and rotation action is supported, multiple volume property actions can be stacked to accomplish complex volume rendering animations. Resultant animation can be saved either as animated in GIF or in MP4 format at selected resolutions suitable for web pages or HDTV applications. The Animator module is built using Slicer’s Sequences and ScreenCapture modules, which can be used to make other types of animation. An example output of the Animator module is provided as an online supplemental material (SOM #4).

## 3. SEGMENTATION AND DATA CLEANUP WORKFLOW

A primary use of the Slicer platform is its volumetric segmentation capabilities. In this paper, we will not delve into specifics of the segmentation process in Slicer, as the best approach to accomplish a segmentation task is typically dataset-dependent and there are many tutorials available both on the Slicer and SlicerMorph websites. We focus here on giving a broad overview of what can be done in Slicer to edit for two different types of 3D data: volumetric and surface representations that depict most of the available 3D specimen images in repositories.

Slicer is not a purpose-specific mesh editing software like the Meshlab. However, there is some support for common mesh editing functionality such as decimation (quadric edge collapse), mirroring, scaling, connectivity, smoothing, boundary detection and others through the Surface Toolbox module. Further operations on meshes, such as clipping with a user defined plane or curve, can be accomplished using the Dynamic Modeler module, a framework for defining parametric surface editing. If the available cleanup and segmentation functionality for 3D models is sufficient for the task at hand, the primary benefit of doing these in Slicer is the preservation of scale and orientation of the model and not risking potential loss of information. Additionally, effects of decimation and other mesh editing on the geometric accuracy of the model can be visualized as a heatmap using the Model to Model Distance extension.

It is also possible to convert a 3D model representation to an image-based segmentation data type at a selected resolution and edit it using the Segment Editor module (Figure 4). In general, converting a surface model representation to a binary volume representation is a lossy operation. If the 3D model at hand was originally derived from a volumetric scan, it is better to use the original volumetric image to re-derive the 3D model. But when that is not available, or the 3D model came from a surface scanner, Segment Editor can be used to remove unwanted structures in the mesh representation of the specimen.

**Figure 4.**
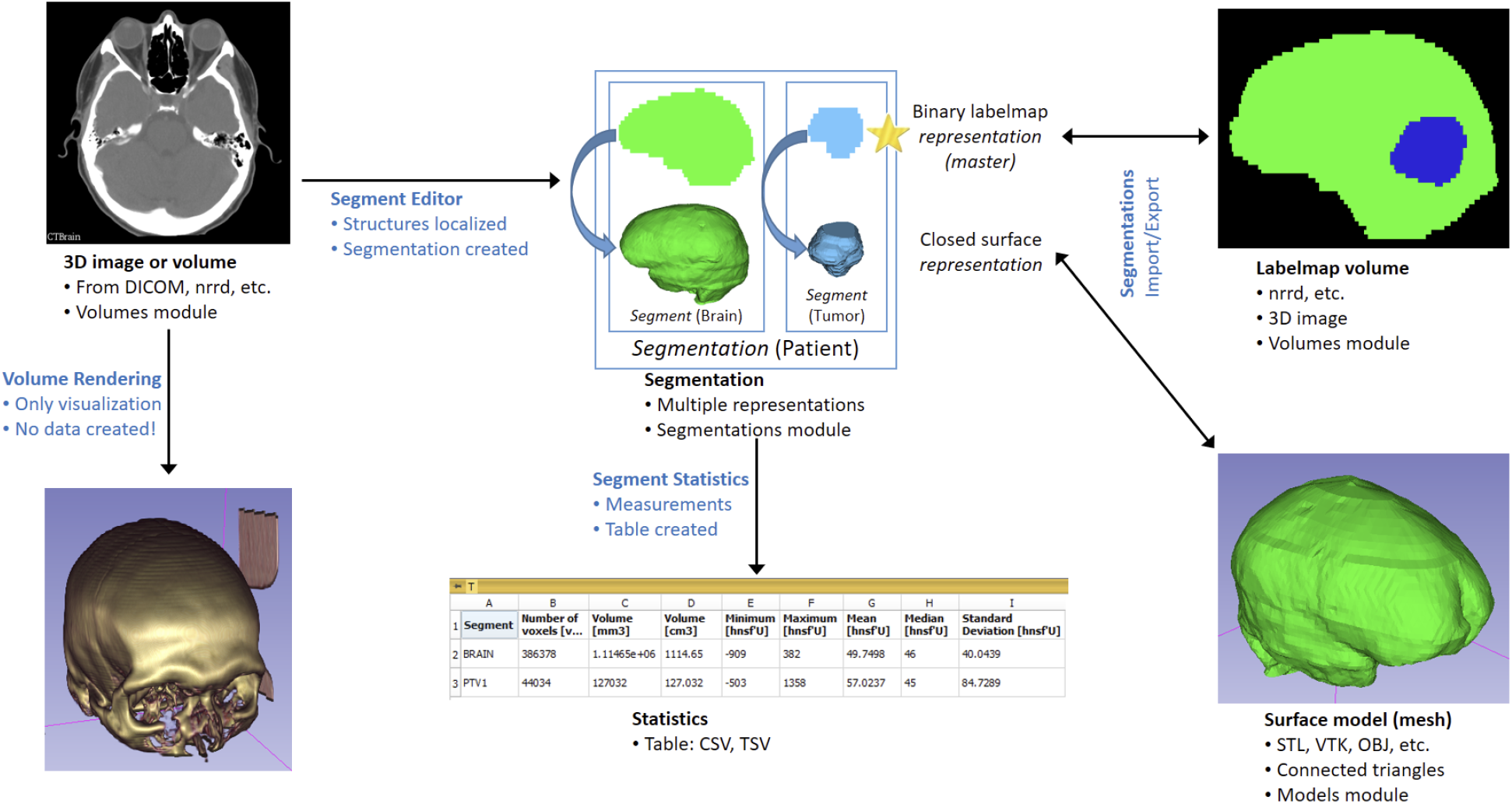
**Relationships of various data types in 3D Slicer** (from 3D Slicer documentation).

As shown in Figure 5, segmentations done by the Segment Editor module can be exported into labelmap or surface model representation. A key difference between segmentation and labelmap representation is that while the former allows for overlapping of segmented structures, the latter does not. For example, in a skull segmentation, a voxel can simultaneously be a member of segments (or classes) skull, mandible, or tooth. If the same segmentation is exported as a labelmap, the same voxel can only be represented as belonging to only one of these classes.

**Figure 5.**
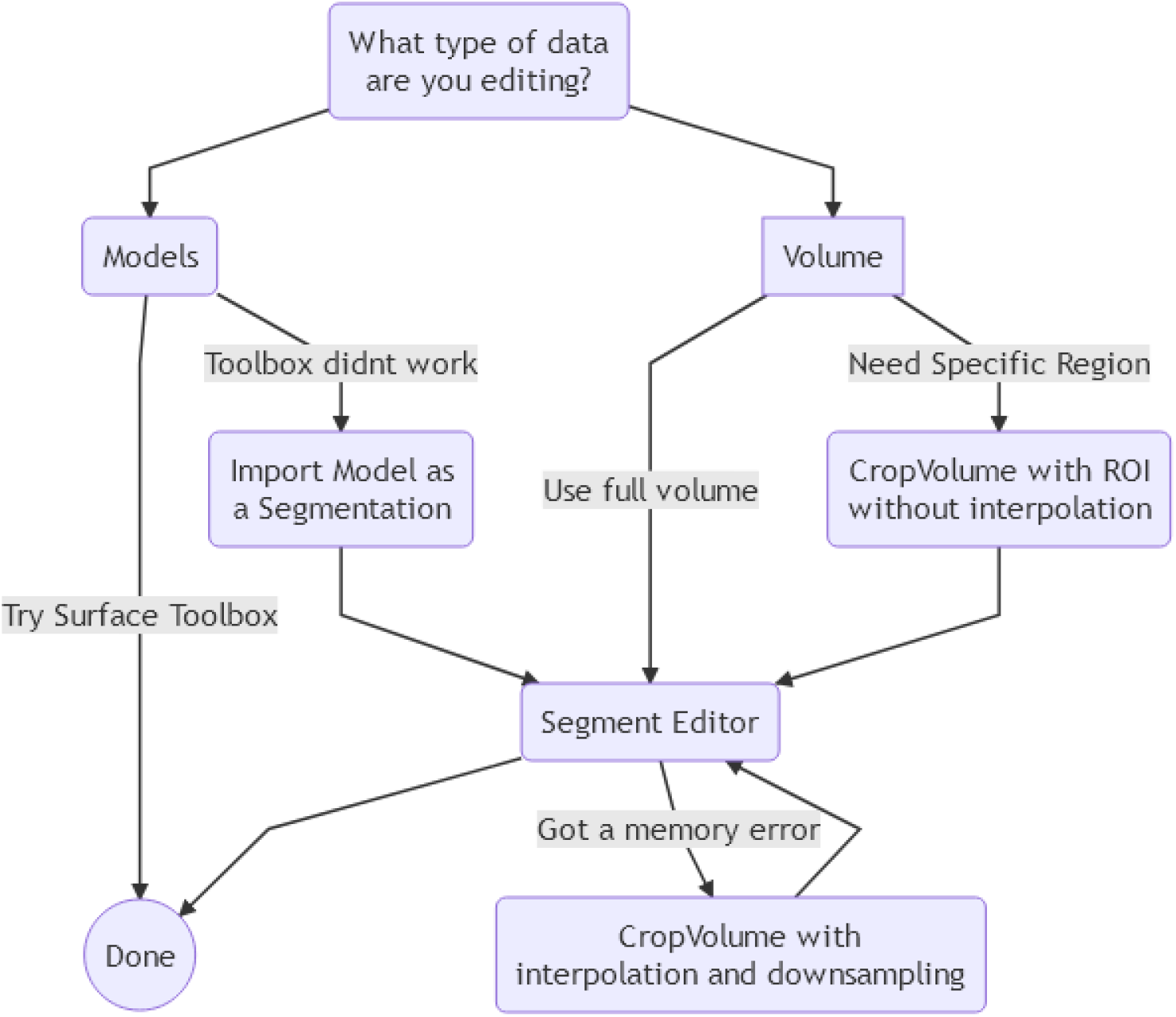
Generic segmentation and data cleanup workflow in 3D Slicer.

Many (but not all) Segment Editor effects (filters) are multi-threaded, and can benefit from running on CPUs with many cores. Running some segmentation effects may transiently require 6-8 times more memory than the size of the volume being segmented to be available during execution. Slicer does not currently support out-of-memory execution, and if the combination of available physical and virtual memory is not sufficient to accommodate this requirement, the operation will be aborted and a Bad Allocation error will result in a warning dialog box suggesting that users save their data and restart the application in case the operation was not cleanly aborted. In such cases, users can either choose to downsample the overall volume or crop the data to a region of interest (ROI). Both of these tasks can be accomplished using the CropVolume module (with and without interpolation).

Finally, we make a plea to the community to refrain using STL and OBJ 3D model formats to share derivative data from their segmentations without careful and explicit notation of the measurement units and coordinate reference (e.g. right-anterior-superior or left-posterior-superior) because this information is essential for proper use of the data. Both these formats suffer from the issue that there is no standardized description of this metadata in the format specification. This can cause scaling and mirroring issues when using these derived datasets across different software with different default units (e.g., millimeters, meters or inches) or spatial assumptions. Like other biomedical visualization software, Slicer supports these formats for historical reasons, and while these formats are sufficient for their original intended use case (i.e., visualization), they are not suitable for quantitative data extraction. We anticipate that emerging open formats (e.g., glTF) will likely to resolve this issue, but meanwhile we advise against the common practice of assuming that all 3D models in STL or OBJ formats are guaranteed to be interoperable, not just for SlicerMorph, but in general. For these reasons, the default save format for model nodes is set to PLY in SlicerMorph customizations.

## 4. MORPHOMETRIC DATA COLLECTION

The Markups module of Slicer provides different types of annotation tools relevant for acquiring morphometric data from specimens. Markup types include fiducials points, lines, angles, open and closed curves, and planes. Among these, fiducials and curves are the most relevant tools for GMM data acquisition. Landmarks belonging to the same class (e.g., skull landmarks from an individual) can be stored in a single fiducial list. Curves, by default are drawn as splines between manually specified control points; although curve type can be changed from spline to linear, polynomial and shortest distance on surface depending on the application. SlicerMorph can resample equidistant points (with or without constraining to the surface of a 3D model) along an arbitrarily drawn curve. This is a core feature of Slicer and is available under the Resample option of Markup module. The Resample option allows one to generate curve-based semi-landmarks from volumes and 3D models (with the optional setting constraining to 3D surface). Default format for the fiducial list is a modified comma-separated flat file format called FCSV. A JSON schema is used for all other Markup types to contain secondary information such as distances or angles between control points, and other derivative measurements. Non-fiducial markups can also be saved in FCSV format, but since this format only allows for saving the control points, it is considered a lossy operation and a warning is indicated in the save dialog box. Alternatively, SlicerMorph-specific ExportAs module can also be used to export these into FCSV format.

Fiducial and curve-based semi-landmarks provide the most common data acquisition method for GMM, i.e., manual annotation. We have supplemented manual digitization with a number of modules that allow generation of patch-based semi-landmarks or pseudo-landmarks from a surface. We use the term semi-landmarks when their generation depends on the user’s input and knowledge of anatomy (e.g., they might be equidistant points along a curve which was initially placed by the user using a number of anatomical landmarks, or they might be derived from a patch bounded by anatomical landmarks). Pseudo-landmarks are geometrically constructed from a model programmatically. No direct anatomical correspondence exists between pseudo-landmarks derived from two different 3D models. Reliance on the user’s knowledge and expertise distinguishes semi-landmarks from pseudo-landmarks.

### Patch-based Semi-landmarks and automation

SlicerMorph’s CreateSemiLMPatches module provides an interface to generate a specified number of semi-landmarks points bounded by fixed anatomical landmarks in the form of equilateral triangular patches. The interface provides a way for the user to experiment to identify the most appropriate set of landmark triplets to span the underlying anatomy. The user selects three bounding landmarks and the sampling rate within the triangular patch. The points from the triangular patch are transferred onto the model surface along a projection vector calculated from the normal vectors at the patch vertices. Two advanced parameters give the user a way to correct cases where the semi-landmarks are projected to an incorrect location. The first parameter, the maximum projection distance, limits the movement of semi-landmark points along the projection vectors. Decreasing the maximum distance is helpful when a model has multiple structures intersecting with a projection vector. The second parameter, normal vector smoothing, adjusts the projection vectors, which are estimated from the normal vectors at the patch vertices. A mesh that is not smooth may produce a projection vector estimate that does not reflect the geometry of the model lying under the patch. Estimating the projection vector from a neighborhood of points around the vertices can improve the result. Increasing the normal vector smoothing parameter increases the size of the neighborhood of points around the vertex points used to estimate the projection vector.

Placement of the triangular grid patches is done iteratively until the region of interest is covered. The user specifies the final set of landmark triplets to construct a fused semi-landmark set, which can be saved as a FCSV file (Figure 6). This process also outputs a landmark connectivity table, which can be used to automatically apply the triangulation pattern to another 3D model with the same set of fixed landmarks, using PlaceSemiLMPatches. In contrast, to the CreateSemiLandmarkPatches module, the PlaceSemiLMPatches module requires the maximum projection distance and the projection vector smoothing to be set to a single value for all patches in the grid. As an alternative to placing the patches on each image independently, the ProjectSemiLMPatches module can be used to transfer semi-landmarks from a template to another 3D model using a thin-plate spline transformation defined by the manual landmark set. This can be used to digitize a dense set of semi-landmark points for a group of samples with a shared manual landmark set

**Figure 6.**
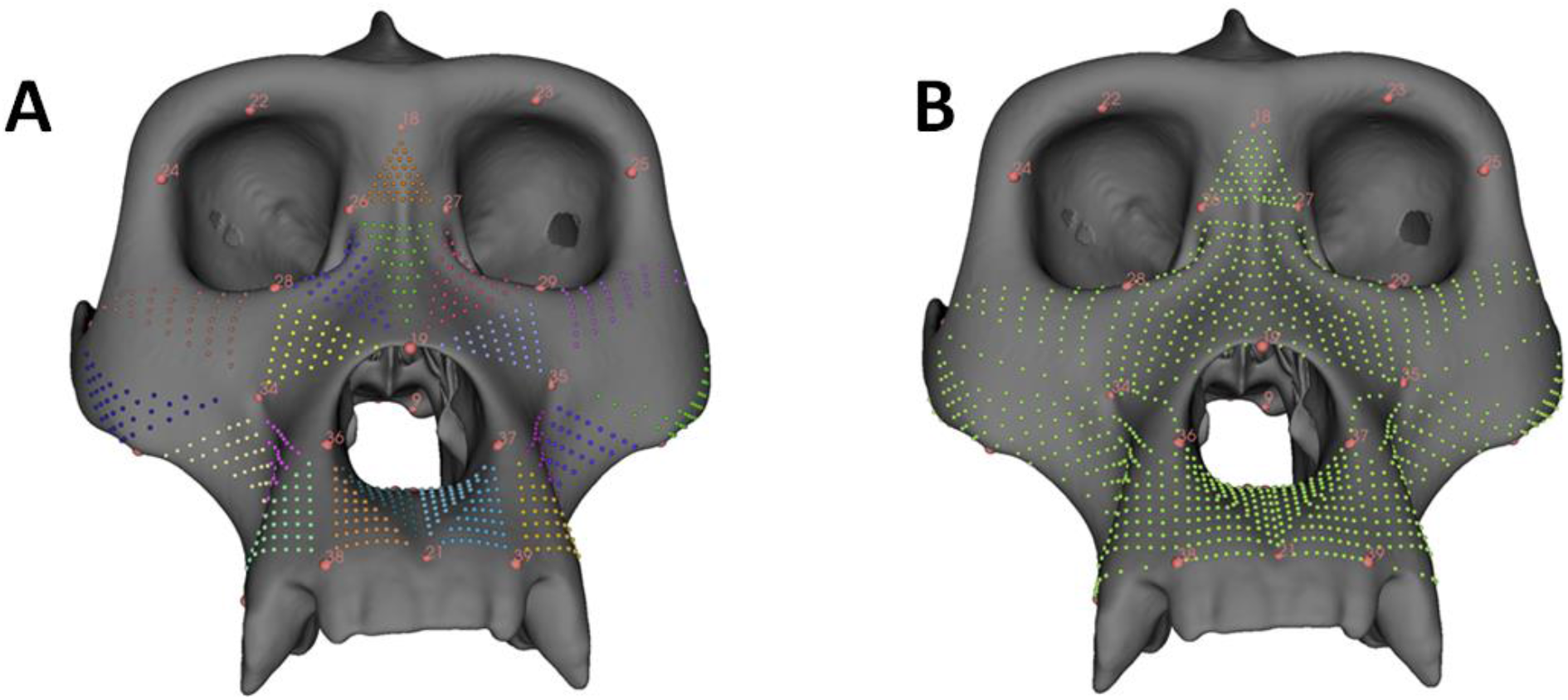
Examples of semi-landmark patches created by the CreateSemiLMPatches module. The module requires the existence of anatomical landmarks (red spheres with numbers) on a surface model. **A.** User specifies triples of these landmarks to create the patches. **B.** Patches are then merged into a single template and the gaps are stitched.

### Pseudo-Landmarks

In genetic studies of organismal shape and form, it is quite common to have large sample sizes of a single organism at a particular developmental time. Placing an initial set of manual landmarks on hundreds or even thousands of individuals, and then using the patch-based semi-landmarking tool is time-consuming. One way to approach this challenge is to use one sample (or if possible, an average of all forms) as a representative, create a dense template of surface points, and transfer them to individual samples. The PseudoLMGenerator module enables the creation of such points, by generating a sparse point representation of the model with the requirement that the points lie on the external surface. Two subsampling methods are supported by this module. In the first, a spatial filter is applied to the model to eliminate points within a user-specified radius, while retaining connectivity. These points are transferred to the most external surface of the mesh by projection along the normal vectors at each point. A second spatial filter is applied to reinforce the spatial constraint after projection to improve regularity of sampling. In the second method, a sphere or ellipse template with regularly sampled points, is selected by the user to approximate the model. The geometric template is placed at the arithmetic center of the model and the points are projected along the normal vectors of the template to the model’s exterior surface. Once the points are placed on the model surface, a second iteration of point projection is performed along the normal vectors of the model. The final spatial filtering step removes points within a user defined radius to enforce regular sampling. The geometric templates provide an improvement in sample regularity when the geometry of a specimen is similar to the selected template shape. Using a sparse point representation of the original geometry provides better sample coverage and regularity for specimens which are not well represented by a sphere or ellipse. The module supports fast and simple comparisons of these approaches to select the most appropriate for a given specimen geometry. If bilateral symmetry needs to be included in the analysis, an optional MarkupsPlane that describes the plane of symmetry can be specified. In this case, only half of the sample will be used to generate a template, and the resultant template will be reflected to the other side. The final set of pseudo-landmarks will indicate the paired landmarks as “normal” and “inverse”, to specify their relationship to the symmetry plane. The landmarks that fall onto the symmetry axis will be indicated as “midline”. It should be noted that most biological specimens are not perfectly bilaterally symmetric and it will be difficult to define a plane of symmetry that will split the specimen into equal left and right halves. If a symmetric template is a requirement, it will be better to symmetrize the selected reference sample prior to pseudo-landmark generation (Figure 7).

**Figure 7.**
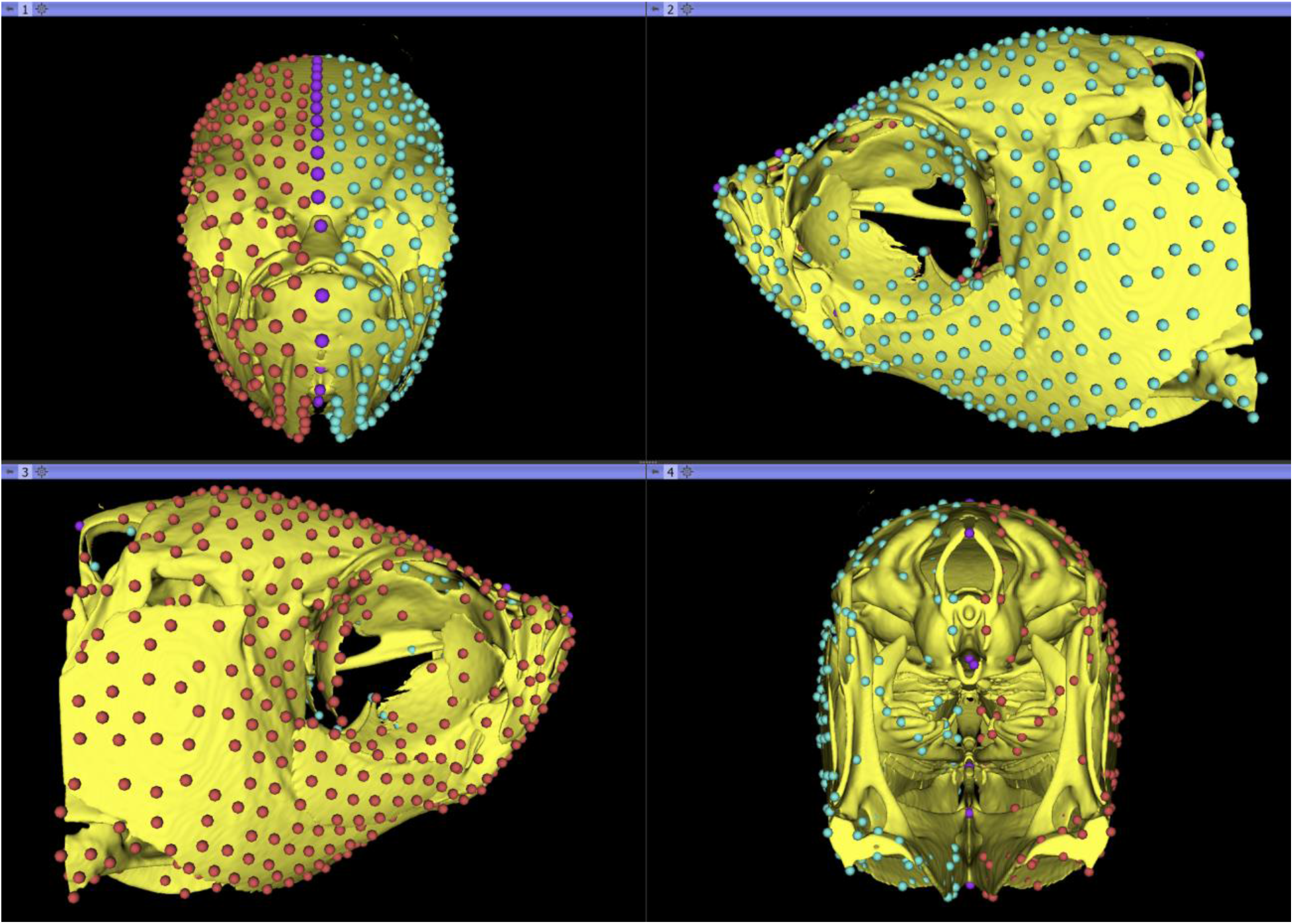
PseudoLMGenerator output on a zebrafish template. This 3D model was created by first generating a synthetic and symmetrized zebrafish template using 23 wild-type adult zebrafish from different clutches and their mirror-reflected copies running through the ANTs multivariate template construction pipeline [78]. The resultant template then was segmented for cranial bones and converted into a 3D model and ran through the PseudoLMGenerator module with the symmetry plane option enabled. Zebrafish dataset was obtained from [67]

As described above, pseudo-landmarks are not anatomically homologous. In fact, if the model that served as the template is replaced with another specimen, a different number of pseudo-landmarks will be acquired and how similar their distribution to the previous attempt will be dependent on how similar the geometries of two models are. The correspondence of the points across samples is usually achieved by using deformable image registration. A volume (or a 3D model) that serves as the reference is deformably registered to the new sample and the landmarks in the template space are transferred to the new subject using this mapping. This is a common approach, particularly in the neuroimaging domain and many different types of linear and deformable registration libraries exist for intensity images and surface models. At least two different registration frameworks, BrainsFit and Elastix, are supported in Slicer. However, deformable registration tasks tend to be computationally intense, and may require large amounts of memory.

### Automated landmarking through point cloud alignment and correspondence analysis (ALPACA)

We developed ALPACA to transfer landmarks from a 3D model to target 3D model(s) through point cloud alignment and deformable 3D model registration. Unlike the PlaceSemiLM module described above, correspondence does not require the presence of a set of fixed landmarks across samples. An optimal set of parameters that gives the best correspondence can be investigated (and outcome can be visualized) in pairwise alignment mode, and then applied to a number of 3D models in batch mode. A paper detailing the specifics of the method and its performance is currently being reviewed, and a preprint version is available on bioRxiv [39]. Below we provide a brief outline of the method and refer the reader to the preprint for technical details of the implementation.

ALPACA uses a lightweight point cloud registration approach to rapidly and accurately register two meshes in 3D space. The procedure has three main steps. As a first step, a reference (source) mesh is isotropically scaled to match a target mesh and both meshes are downsampled into point clouds. After downsampling, the two point clouds are rigidly aligned using a combination of a feature-based global registration algorithm (FPFH-based RANSAC; [20]) and a local registration algorithm (Point-to-plane ICP; [40]). As a final step, the two rigidly aligned point clouds are then subject to a deformable registration step (CPD; [41]) and the warped source model landmarks are then transferred to the target mesh. In other words, ALPACA performs automated landmarking by transferring the landmark position of a single template into other specimens (Figure 8).

**Figure 8.**
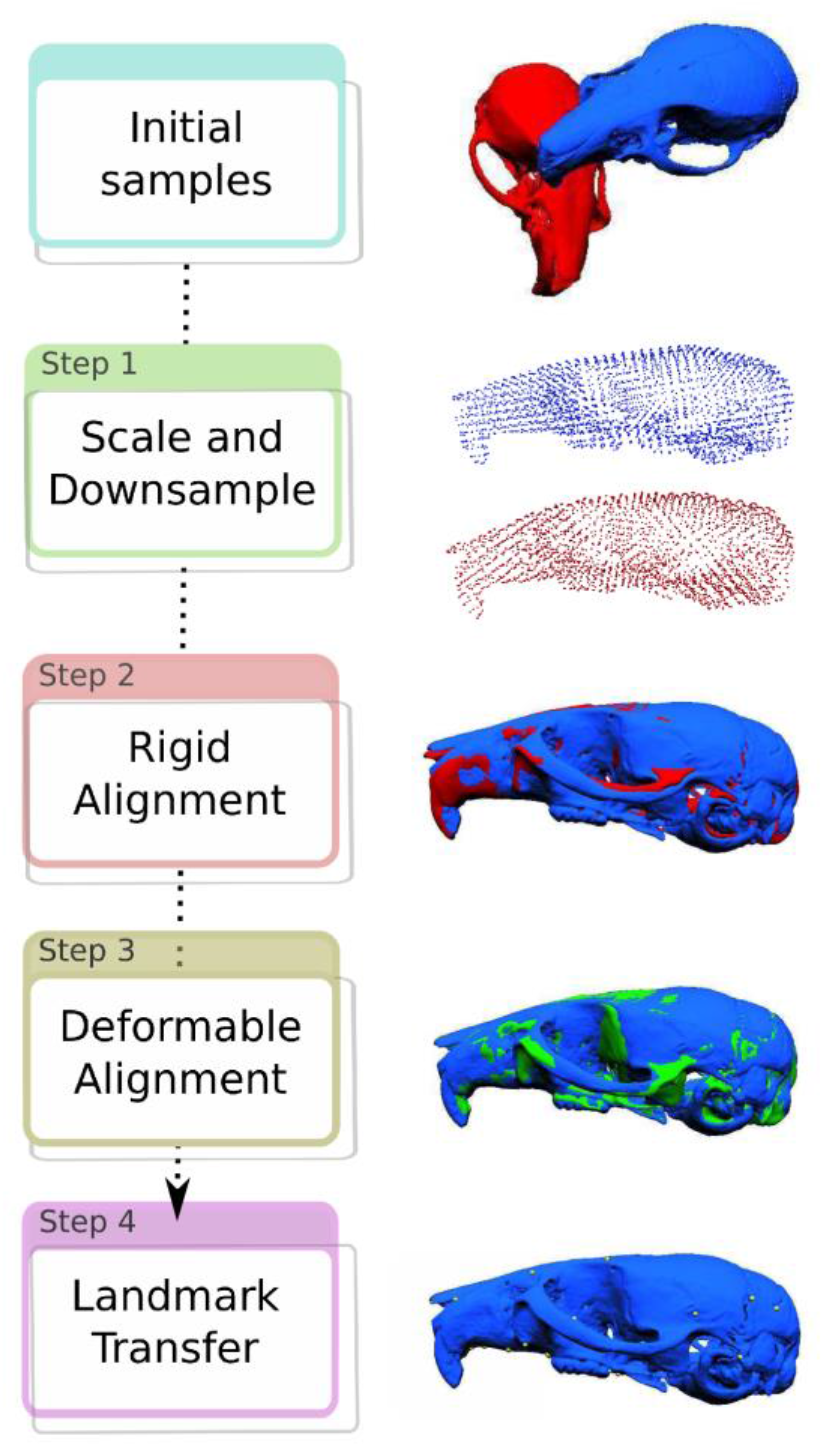
**ALPACA** module conceptual workflow.

The main benefits of ALPACA are that it is a general algorithm that should work on any biological structure of interest, and it is also intuitive, allowing users with no prior programming experience to perform automated landmarking in their own systems. The ALPACA panel contains two different modes of functionality: a pairwise tab and a batch-processing tab. Users should use the pairwise tab to search for the best combination of registration parameters for their dataset. Once the results are satisfactory, users can then apply the procedure to a larger array of samples in batch mode. Note that ALPACA was developed to work well with most biological structures, so all tunable parameters are located under the ‘Advanced parameter settings’ tab. In most practical applications, the main parameters that will need to be tuned are the deformable registration parameters, as those can vary considerably across structures and/or species. A preprint detailing the specifics on the ALPACA implementation and its performance on a number of mammalian skulls can be found in bioRxiv [39].

### Auto3Dgm

Is a Python3 implementation of a previously developed “automated 3D geometric morphometrics” (auto3dgm) method that aims to align 3D mesh or point cloud representations of anatomical structures [20]. The method is perhaps most useful for samples in which the shapes of the structures are quite dissimilar, or for which a very few of anatomically homologous points can be reliably identified [42,43]. In samples with such morphologically diverse and disparate shapes, it can be challenging to use a single specimen as a representative for annotating semi- or pseudo-landmarks with the modules described above. In theory, the more species diversity in a study sample, the more disparity will accumulate. Thus, it is not surprising that most of the studies published using auto3dgm focus on macroevolutionary questions with multiple species in each sample [36,44–53].

The method has even been employed for analysis of non-anatomical data [54,55]. However, it can still be powerful and preferable in certain monospecific samples [56]. A key point that users should consider is that the automatic pseudo-landmarks generated by auto3dgm cannot be expected to preserve intuitive notions of “homology” between shapes with very different proportions. The team that built this module maintains a website with instructions, downloads and example datasets at https://toothandclaw.github.io/. Much of the practical information presented at the end of this section is described in more detail on the website and existing literature. Here, we provide a broad overview of the Python implementation in Slicer.

In the SlicerMorph implementation of auto3dgm, two analytical steps must be completed sequentially. First, shapes are downsampled to a target number of pseudo-landmarks. Pseudo-landmarks can be “evenly” distributed according to different criteria including farthest point sampling, geodesic distance or minimization uncertainty using a Gaussian process model [57]. Other approaches like using geodesic distance or minimizing bending energy are not currently supported. The more pseudo-landmarks are created, the more stable the representation of shape similarities and differences in the target sample [20,36]. However, the computational complexity also increases with increased number of pseudo-landmarks. Most studies use 500-1,500 pseudo-landmarks per object. Using more feature aware methods of landmark placement, like Gaussian process models, may increase the stability and information content of selected landmarks allowing stable shape representations with fewer landmarks (probably by an order of magnitude). At this stage the software checks for and reports on any “bad” meshes that will cause the analysis to fail, so they can be cleaned or removed.

Next, alignment of shapes in the sample is computed using a Procrustes distance matrix reflecting pairwise alignments and the minimum spanning tree among shapes implied by that matrix. Pairwise alignments are computed using the iterative closest points (ICP) algorithm [58], which searches for the best rigid transformation and the best match of points that produce the least Procrustes distance between two shapes. This method tends to be very slow and computationally expensive, so we apply certain assumptions to help find the optimal alignment more quickly. In auto3dgm, we provide several standard kinds of rigid transformations as initial seeds for ICP. These standards are defined such that we expect one of them to be very close to the optimal alignment in many cases. For each initial rigid transformation, ICP returns a Procrustes distance between the shapes (which are standardized to a centroid of 1.0 prior to comparison) by identifying the closest points in the shape and continuing with additional iterations of adjusting point correspondence and rigid transformation. The alignment that results in the smallest Procrustes distance out of the eight is used for the next steps while the other seven are discarded.

The above process is repeated for all pairs of shapes and results in the Procrustes distance matrix of individual pairwise distances. The assumption that any of three principal axes of variation between two shapes are anatomically homologous is often critically flawed when the two shapes are very different, however the next phase of the algorithm is designed to exclude all the incorrect pairwise alignments. In this step, the pairwise distances are used to construct a minimum spanning tree, which is the graph that connects all the cases in the sample using the smallest sum distance. This graph is used as a path for determining correspondence and alignment between dissimilar objects. If two objects are dissimilar, in most cases the Procrustes distance separating them will be quite large and it will be excluded from the minimum spanning tree [59]. Using this approach the algorithm can achieve biologically meaningful alignment and correspondence in samples that include very dissimilar shapes as long as there is also a good sampling of intermediate shapes.

In practice, the process of determining alignment and correspondence is run twice, once on a low-density point subsample and once on a high-density sample. The analysis on the low-density sample is much faster and establishes roughly correct correspondences. This saves time in the pairwise alignment step during the high-density stage by giving ICP a single initial guess of rigid transformation from which ICP is more likely to return the optimal alignment.

Once the auto3dgm computations are completed, it is possible to visualize the resultant pseudo-landmark distribution and model alignment (Figure 9). Auto3dgm yields several outputs. Of primary interest to most users will be a series of FCSV files that represent the identities and coordinates of pseudo-landmarks generated by the analysis. There will be a set of FCSV files for both the low- and high-density passes. For both passes, two versions of each set of FCSV dataset are produced. In one set, the pseudo-landmarks coordinates represent the final aligned and scaled configuration producing the procrustes distances used for the minimum spanning tree. In the other, these pseudo-landmarks have been back transformed to their Original Shape Space (OSS). While either of these data can be used in Slicermorph geometric morphometric analyses of shape covariation, we advise to use the data in the Original Shape Space to preserve the original scale of the data. GPA will need to be rerun as an initial step with the OSS fcsv output. Additionally the program outputs a copy of the input meshes re-aligned to each other using pseudo-landmark correspondences, which may be useful for certain kinds of measurements [60] or as starting points for other algorithmic workflows [61], or simply to visualize the resultant alignment in SlicerMorph.

**Figure 9.**
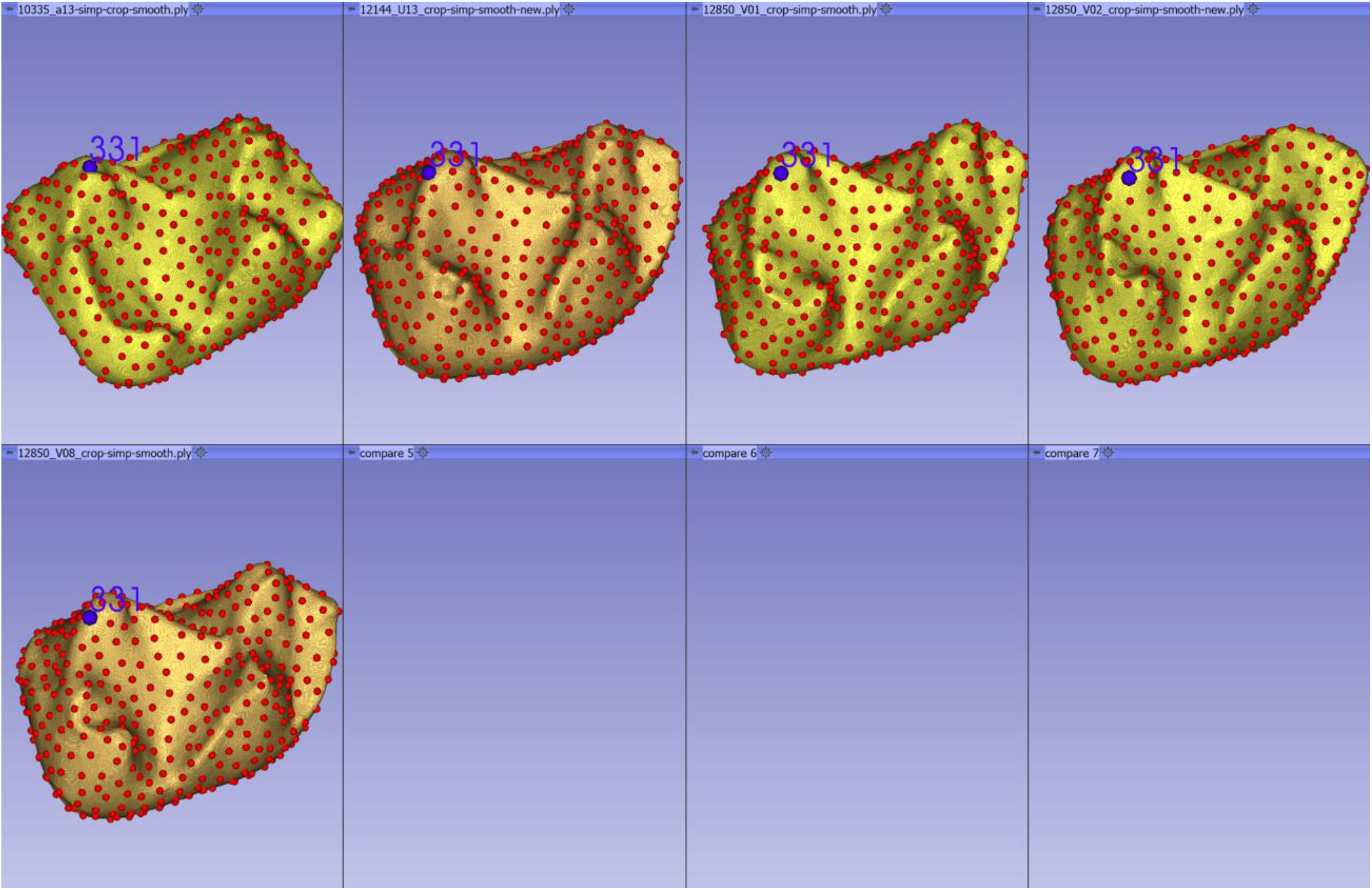
Auto3dgm output. Users can visualize the alignment and point correspondence calculated by auto3gm run using the visualization tab of the module. Here, five molars from the sample data provided by auto3dgm module were aligned using 400 points and 1000 iterations. Users can interactively investigate whether the calculated point correspondences are sufficiently good for their analysis using the available display options of the Markups module. In practice, using larger sample sizes with higher morphological disparity and higher point numbers result in better correspondences across samples.

### Considerations about choosing a Geometric Morphometrics data acquisition workflow

Of the workflows we present, GMM data collection is the most complex to describe due to the variety in study designs, modalities and conflicting requirements on accuracy, repeatability, throughput and generalizability (Figure 10). Additionally, since GMM are used to address a wide-range of biological questions from developmental biology to comparative phylogenetics, the nature of datasets and the phenotypic variability they represent can be very different. An evo/devo study on brain shape and evolution in mammals may have hundreds of different species, in which each species is represented by a couple of samples. On the other hand, a quantitative genetics study of shape may require hundreds to thousands of individuals of the same species at a very precisely timed developmental stage or from a single geographical locality. A study on developmental origins of organismal form may have a large sample size representing multiple developmental stages, each of which can be very different from the others, even though they are from the same species. While data collection for all of these studies can be accomplished using the tools in SlicerMorph, they require different approaches. For example, recent automated data acquisition can increase the analytical throughput of morphometric studies so large sample sizes can be analyzed [20–23,25,62]. But most, if not all, automated methods require pre-processing of the 3D data. Most often, whichever anatomical region is going to be analyzed, it needs to be segmented carefully so identical structures exist in all samples. Hence, if this preprocessing step cannot be automated, there will still be a time (and therefore sample) limiting step, even though the landmark collection is automated. On the other hand, manual landmarking, particularly in Slicer can be surprisingly time-effective. In fact, once the user is comfortable with the software and familiar with the anatomy, annotation of landmarks from a single specimen (e.g., standard set of craniometric landmarks on a mammalian skull) is usually a matter of minutes, even when starting from an image sequence of an articulated vertebrate skeleton. So, if a set of manual landmarks is sufficient to answer the biological problem, manual landmarking can be an effective and quick solution even for sample sizes in the hundreds, for example in the case of that hypothetical study on brain shape and evolution in mammals.

**Figure 10.**
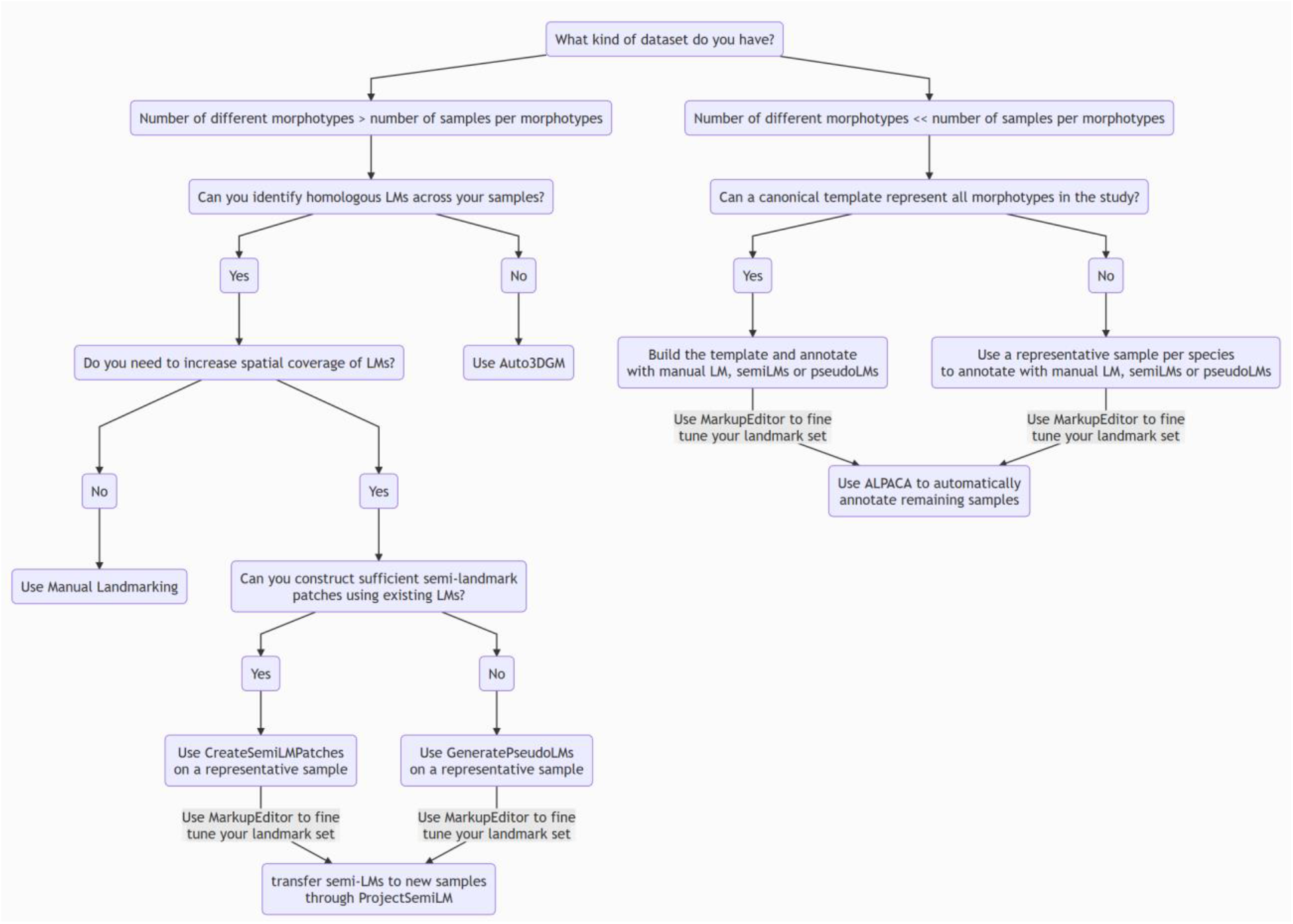
Geometric morphometrics data collection workflow

As noted above, the main drawback of using manual landmarks for GMM is not the pace of data acquisition, but the concern for repeatability due to well-documented intra and inter-observer biases. So while conducting manual landmarking users should be mindful of the impact of these biases to the particular question they are trying to answer. Unfortunately most of the time the only way to capture these biases is through repeated landmarking of the same dataset by multiple experts [63,64]. There might be practical limits to manual landmarking, such as an inadequate number of anatomical landmarks to represent the topology of the anatomical region in sufficient complexity. A typical example of such structure is the ankle and wrist bones of the mammals, or any bony appendage that lacks prominent anatomical markers (projections, sutures etc). In this case, some sort of computational method of establishing correspondence and generating additional landmarks across samples will be necessary.

Users must consider how certain algorithms work, and how that may impact their data collection efforts. For example, while auto3Dgm can deal with dissimilar shapes, and can establish both alignment and correspondence of numerous pseudo-landmarks, that correspondence is unique to the samples included in the analysis. It is not possible to retrospectively add a couple more samples to an existing analysis as they become available, because the correspondence across samples has to be re-established. This is in contrast to deformable registration based methods (e.g., ALPACA), in which it is possible to re-use the reference model and its associated landmark set to annotate a new sample at any time. Methods also differ in their computational costs and expectation from the users. Of the automated methods in SlicerMorph, auto3Dgm is much more computationally taxing, whereas ALPACA is fairly lightweight for similar datasets. That’s primarily due to ALPACA being a supervised method. While both modules take a set of 3D models to be landmarked and output a set of landmarks on each of these models, ALPACA requires that a reference template of landmarks and a 3D model exist, while auto3Dgm not only aligns the models but also creates the pseudo-landmarks.

Scans of damaged or partial specimens can be manually landmarked, and if necessary, certain landmarks can be skipped and dropped from the analysis to retain these samples in the analysis. On the other hand, most automated methods, including ALPACA and auto3Dgm, require complete specimens to be included in the analysis. Thus, there is currently no one method that fulfills the conflicting requirements of accuracy, repeatability, throughput and while still being broadly applicable to all types of GMM inquiries. That’s the primary reason we have implemented a number of GMM data collection approaches in SlicerMorph, but undoubtedly more methods will be developed, and Slicer provides an excellent platform for this future development. We advise users to follow our tutorials to get familiar with what each module accomplishes and where they might benefit in using that. Here, we also provided a workflow for landmarking that tries to find a balance between time cost of GMM data acquisition and the phenotypic variability in the dataset (Figure 10). To do that, we introduce the term ‘morphotype’ for a concept to aid our presentation of workflow options. We define morphotypes as groups of 3D data that are expected to be phenotypically more similar to each other than they are to other morphotypes. So, depending on the study, a morphotype can be a developmental time of a single organism, a population from a single geographical locality, a species, or other taxonomic levels. In general, it is usually easier and time-efficient to implement a workflow that will be repeatable, high-throughput and automated within a single “morphotype” [23,65–67]. But again, the workflow we present should be taken as a general advice more so than finding a module that will perform a specific function. Ultimately, it will be up to investigators to find the most appropriate workflow that will help them address their particular question (Figure 10).

MarkupEditor is a SlicerMorph-specific add-on to interact with dense landmark sets associated with a model. It allows for interactive selection of visible landmarks by drawing a 3D curve to define subsets. The functionality is available by hovering the mouse over any fiducial point and right-clicking on it to show the context menu. After drawing the curve, 1) the points inside the curve can be set to ‘selected’ state, and outside to ‘unselected’ state, which is indicated by a different color then the original, 2) points inside the curve can be added to the current selection, or 3) points inside the curve can be removed from the current selection. Once the selection is finalized, the user can employ the Markup modules Control Point tab to operate on these landmarks from the existing landmark set. For example, this would be a situation where the PseudoLMGenerator is used to generate pseudo-landmarks across the entire surface of a 3D skull model, but the user wants to retain landmarks only from a specific-region (e.g., facial skeleton, or neurocranium). Alternatively, users can copy/cut and paste the selected subset as a new fiducial list, or simply obtain the landmark indices of the subset to use in the downstream analyses (e.g., as modules in an integration and modularity analysis).

Whether they are manual landmarks, semi-landmarks or pseudo-landmarks, all GMM data modules in SlicerMorph will output the results in FCSV format. As noted above, users can perform format conversion from Morphologika and IDAV Landmark Editor to FCSV using the included convenience modules.

## 5. GENERALIZED PROCRUSTES ANALYSIS (GPA)

The primary goal of the SlicerMorph’s GPA module is to provide the user to vet their data collection efforts in the same platform that is used for digitization. This way, users can check for digitization errors, identify outliers in their dataset and -if necessary-fix issues within Slicer without having to move data back and forth between different analytical software. To initiate the GPA, the user needs to provide a folder of FCSV files where each specimen is identified by the filename. A time-stamped output folder is created under the specified output folder. By default, all landmark configurations are scaled to unit size to conduct GPA, but this step can be optionally skipped to preserve the scale of the data. It is possible to specify the landmarks to exclude, but not the samples. Should users want to exclude specimens, they need to remove those FCSV files from the folder. When executed, the GPA module will align the landmarks and apply principal component analysis (PCA) to the procrustes residuals. Outputs for procrustes aligned coordinates, centroid size, PC scores, and coefficients of individual eigenvectors and the variance associated with eigenvalues are saved as separate files in csv format in the output folder.

As the first step of data vetting, we suggest users to quickly review the Procrustes Distances plot and table to identify potential outliers. Likewise, plotting Procrustes Distance variances associated with landmarks as spheres (average of three axes) or ellipses (each axis calculated independently) can give visual clues about what landmark(s) might be problematic (Figure 11A-C). Due to the way GPA finds an optimum solution that minimizes total Procrustes Distance, it is not meaningful to interpret each landmark variance on its own. Users should refrain from attributing any direct interpretation of these variance plots, but use them strictly for diagnosis of potentially problematic landmarks that might be indicative of digitization problems. Large amount of variation in a landmark relative to others may be an indication of a number of non-biological causes of variation such as wrong ordering of landmarks in some samples, a change in the way how these landmarks are recorded (aka learner’s bias), anatomical structure not being clearly visible in some of the sample, etc. Such errors should be minimized. Plots of mean shapes (with or without a reference model) can be used to investigate the consensus landmark configuration (Figure 11A).

**Figure 11.**
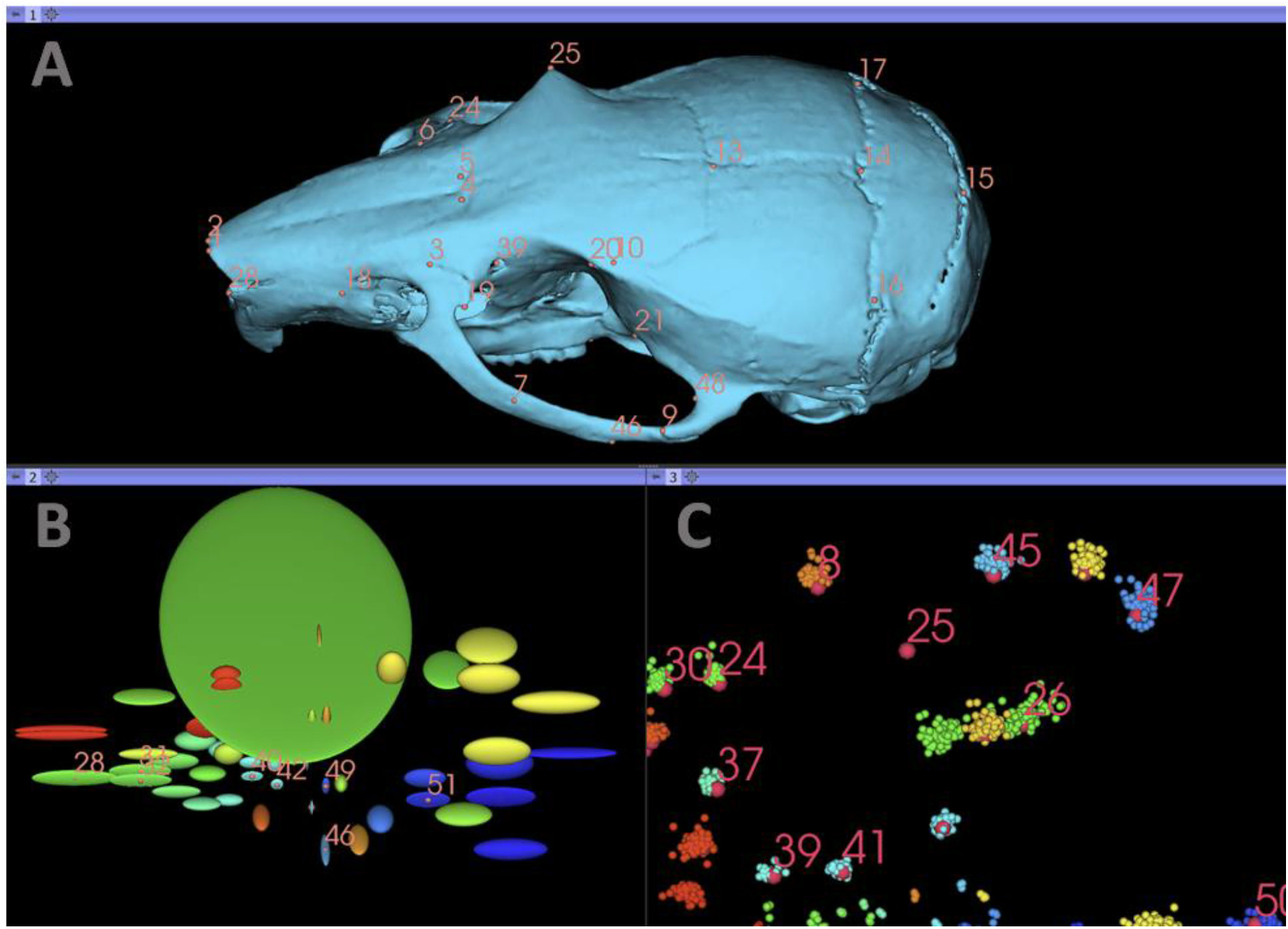
Visualizing problematic landmarks via GPA module. This figure was generated using one of the sample dataset distributed by SlicerMorph. Dataset contained coordinates of 55 landmarks from 126 mouse skulls. The figure shows three potential ways of visualizing and identifying problematic landmarks (landmark 25 in this example) in a dataset. **A.** Selected reference sample is warped into mean shape. **B.** Visualization of relative landmark variances in the dataset. In this case, variances for each axis is calculated independently, resulting in ellipsoidal shapes centered on the mean coordinate of the landmark. **C.** Visualization of GPA aligned coordinates as point clouds (smaller spheres) and mean coordinates. Image was zoomed into the region of landmark 25. The green cluster of points in the center are the procrustes aligned coordinates of landmark 25 for most of the samples, but a large error in a few samples causes a large shift in the mean coordinate for landmark 25. In all three, light red text and spheres indicates the position of the mean shape coordinates and the landmark label.

To understand shape variation trends in the data and ordination of the samples in the multivariate morphospace, users can generate bivariate plots of individual PC scores and use the static and interactive visualization tools in GPA module (Figure 12). Optionally a categorical grouping variable can be added within SlicerMorph to label data points using the table. There are two ways to visualize eigenvectors associated with each PC. A 3D static representation can be achieved by using the lollipop plot option, in which each eigenvector is represented as a line moving away from the mean shape coordinates, resembling lollipops (Figure 12A). Alternatively, to create an interactive representation of shape variability associated with each eigenvector, users can opt to use only mean shape configuration, or a reference model. In case of landmark visualization, mean (consensus shape) is used to visualize the results. If a 3D model and its set of landmarks are chosen as the reference to use for 3D visualization (Figure 12F), the selected model is first deformed into the mean shape using TPS as the initial step, and then eigenvectors from selected PCs are used to further deform the model (Figure 12G). The scaling coefficient associated with each PC is arbitrary and aims to provide a noticeable deformation for visualization purposes. Finally, it is possible to capture this dynamic visualization as an animation and save it to disk either as an animated GIF or in MP4 format. Further stylistic customizations of plots, landmark points and models can be changed using the appropriate module’s (e.g., Plots, Markups, Models etc) display properties. Should the user prefer to visualize the mean and warped 3D model in the same 3D viewer, default GPA window layout can be changed to different layouts (e.g., 3D only, Plot only, Table only) using the layout manager and the display properties of the Model module can be adjusted accordingly.

**Figure 12.**
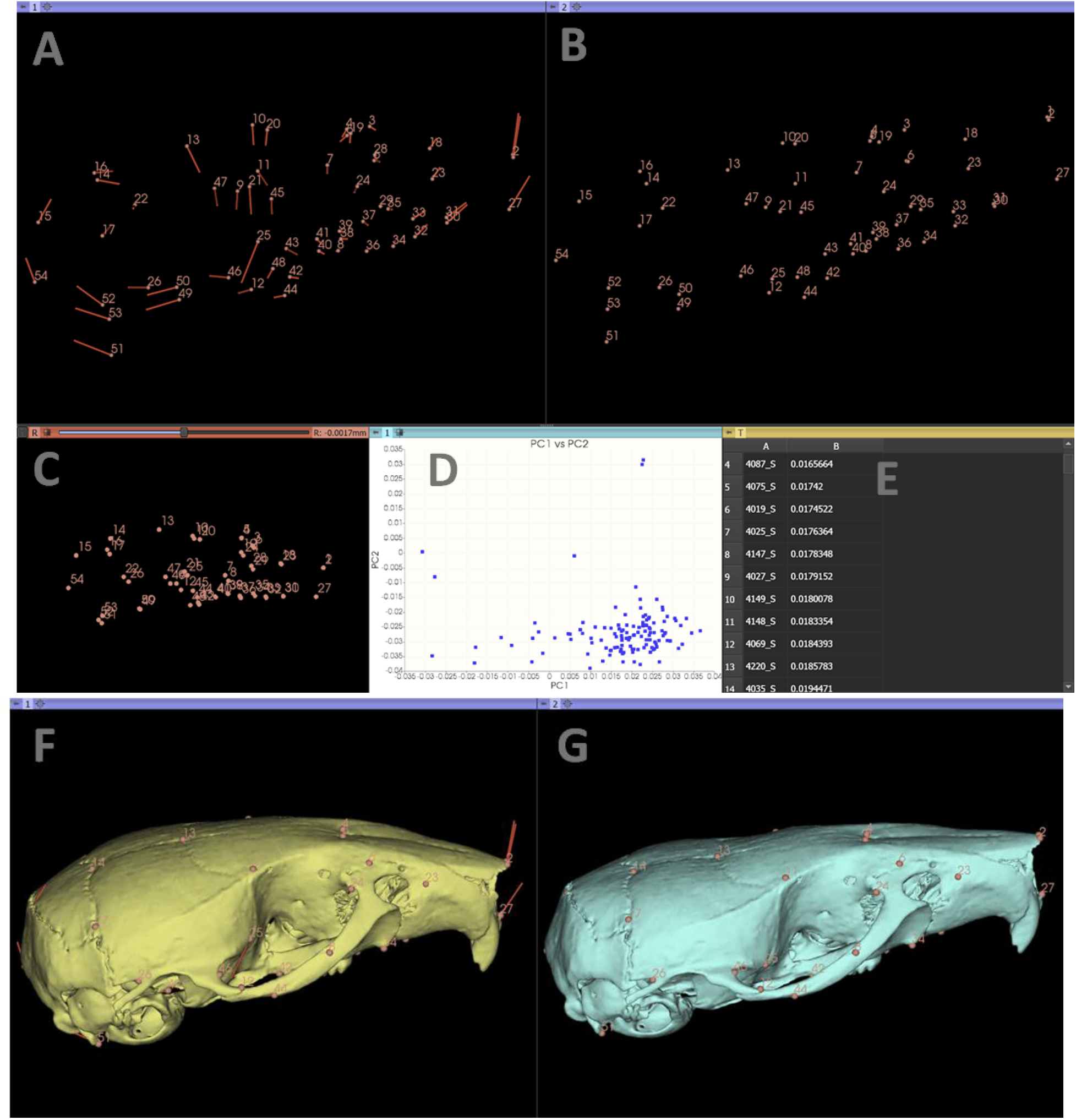
Visualizing Shape Variance. GPA module has its own layout which is initially populated by a table of two 3D viewers (**A, B**), 2D projection of mean shape (**C**), a plot tab (**D**), and a table tab that displays sample IDs, procrustes distances and -if provided-other covariates (**E**). In addition to displaying mean shapes in 3D, if selected, eigenvectors associated with selected PC can also be visualized in the left 3D viewer (**A).** Right 3D viewer shows the new coordinates of the landmarks after PC deformation is applied (**B**). The scaling coefficient is arbitrary to provide a visual effect of displacement of points. Alternatively, users can choose one of their samples as a reference model, which is first warped into mean shape (**F**), and then PC deformation is applied (**G**).

Currently, GPA module is intended to function primarily as a data exploration and visualization tool. As such, it does not feature any statistical shape analysis capabilities such as Procrustes Anova [68], symmetry decomposition [69], multivariate allometry [70]). R statistical language is rather rich with different statistical shape analysis libraries [71–75], and we encourage the users to explore these libraries for complex analyses such as shape models, integration of GMM with phylogenetic comparative methods, modularity analysis and others, using the outputs from the GPA module. A brief example of importing GPA output into R to be analyze with R/geomorph package is provided as a supplementary material (SOM #5).

## CONCLUSION

We have developed the SlicerMorph extension in response to challenges we, as well as our collaborators and colleagues, face in studying morphological variability using 3D data. Our goal was to establish 3D Slicer as a one-stop platform to do most (if not all) the tasks associated with the study of digital morphology. As it stands, all common tasks (data conversion, downsampling, segmentation, visualization, taking measurements, landmarking, shape variation decomposition, and animations) of digital morphology can be accomplished within Slicer using SlicerMorph. More advanced geometric morphometric analysis such as building and testing statistical shape models and symmetry analyses can be implemented using the built-in Python interpreter and available libraries from the python ecosystem; or data from SlicerMorph can be easily imported in R (SOM #4).

Another benefit of using a single platform for digital quantitative morphology is that researchers now can investigate how all the preprocessing steps (import, segmentation, surface generation, downsampling) to generate the geometric model for shape analysis can impact the downstream analysis more conveniently. While errors associated with the landmark digitization and how those may affect the shape analyses have seen a substantial treatment in literature [21,63,64,76,77], not much has been done to investigate how various methods of image processing affect the 3D model generation, and subsequently the data collection This is mostly due to the fragmented workflows that require serial use of different software and difficulty of measuring the impact of changing a single parameter on the final outcome.

In the present paper, we provided a general overview of how common workflows associated with digital morphology can be accomplished with SlicerMorph. We chose not to give explicit, step-by-step instructions because as a research software driven by the needs of its large community both Slicer and SlicerMorph constantly evolve. While individual modules and their functionality (e.g., CropVolume) continue to exist, how users interact with them may change due to revised or added functionality in future; rendering a step-by-step tutorial in a publication like this out-of-date in a short amount of time. Instead, online tutorials are updated frequently and can be kept up-to-date.

In future, we plan to create a seamless ‘bridge’ between SlicerMorph and R statistical language so that such complex geometric morphometric analyses can still be done in R, but visualized in SlicerMorph interactively. Slicer’s plotting infrastructure also allows for interaction with data points (such as selection, drawing) which can be used to create a more interactive exploration of the morphospace. We also anticipate deploying Slicer and SlicerMorph on research and commercial cloud computing platforms, for users who need to have access to more computing power to deal with larger and larger datasets, or make sure their students and trainees have consistent experience while teaching a course or a workshop.

We hope that the tools we developed for SlicerMorph will be useful to build a community of organismal biologists and quantitative morphologists that value open science and enable easier collaboration.

## ACKNOWLEDGEMENT

SlicerMorph development and short courses are funded by NSF Advances in Biological Informatics grants to AMM (1759883), AS (1759637) and DM (1759839). We would like to thank the members of our advisory committee, Dean Adams, Anjali Goswami, David Polly and Jim Rohlf, for their guidance and input into the project. Finally, we like to acknowledge the support and contribution of the 3D Slicer developer community to the development of SlicerMorph in general and Dr. Andras Lasso’s (Queen’s University) mentorship in particular. We also would like to acknowledge Dr. Ronald Kwon (University of Washington), who gave access to the zebrafish dataset used to generate Figure 7.

Authors declare no conflict of interest.

## AUTHOR CONTRIBUTIONS

AMM conceived the project, acquired the funding, oversaw the design, created documentation and wrote the initial manuscript. SR led the software development. SP, AP, JW contributed modules to SlicerMorph. KD, DB and AS contributed to the module designs and documentation.

All authors contributed to editing of the manuscript and agreed to the final form of the paper.

## SUPPLEMENTAL ONLINE MATERIALS

### SOM 1. INSTRUCTIONS FOR OBTAINING SLICER AND SLICERMORPH

1. Slicer can be downloaded from https://download.slicer.org. The current stable version (4.11.20200930, r29402) should be used, as SlicerMorph is not available for older versions (v4.10 and earlier) and preview versions may be unstable. Hardware requirements, and operating system specific instructions for installing Slicer can be found at: https://slicer.readthedocs.io/en/latest/user_guide/getting_started.html#
2. Once Slicer is installed, open the Extension Manager and search for SlicerMorph extension. Click install, and wait all additional dependencies are installed and restart Slicer. After the install SlicerMorph will be listed as a top-level folder in the Module selection toolbar with modules organized into three subfolders (Input Output, Geometric Morphometrics and Utilities).

### SOM 2. LEARNING MORE ABOUT 3D SLICER AND SLICERMORPH

Official documentation for Slicer can be found at https://slicer.readthedocs.io.

Official user guide provides the background for core features such as user interface elements, mouse and keyboard operations, loading and importing data, visualization and segmentation.

Information about SlicerMorph specific modules can be found at SlicerMorph’s main github repository, https://github.com/SlicerMorph/SlicerMorph. Clicking on the name of the module will take the user to the documentation page specific to the module. Module documentation page also provides a link to the in-depth tutorials for the specific module or the task. Link to the module documentation is also provided in the Slicer’s inline Help tab that is available as the top section of any active module.

Content from our short courses are publicly available on SlicerMorph github page. The latest incantation can be found at https://github.com/SlicerMorph/S_2020

### SOM 3. GLOSSARY OF TERMS

#### Anatomical landmark

A term that refers to both Type I (juxtaposition of anatomical structures; e.g., suture) or Type II (extremities of anatomical structures) landmarks of Bookstein.

#### Markups

A category that includes point lists, lines, angles, curves, and planes all defined by a set of control points.

#### Medical Reality Markup Language (MRML)

Is a data model developed to represent all data sets that may be used in biomedical software applications [79]. When an MRML data is saved, an XML document is created with .mrml file extension. This file contains an index of all data sets in the scene and it may refer to other data files for bulk data storage. A .mrb file is a zip archive containing a .mrml file and a data directory with all the referenced bulk data. More specific information about MRML, MRML Scene and MRML Nodes and data representation in Slicer can be found at https://slicer.readthedocs.io/en/latest/developer_guide/mrml_overview.html#mrml-overview.

#### Model

Refers to both surface (such as an PLY/STL/OBJ file), and volumetric (such as a tetrahedral mesh for finite element analysis) polygon mesh.

#### MRML Node

All objects loaded into (and generated by) Slicer are represented as MRML nodes that are visible under the Data module (All tab). Different nodes have different properties associated with them that are set by different modules. Right-clicking on a node and choosing Edit Properties action will take the user to the module that can set those parameters. Each MRML node has a unique ID in the scene, has a name, custom attributes (key:value pairs), and a number of additional properties to store information specific to its data type. Node types include image volume, surface mesh, point set, transformation, etc. Nodes emit events when their data changes so that different parts of the user interface, such as tables or rendered views can be updated to reflect the current state. In this way the various elements of the program are kept in sync when node data is modified by user actions, scripts, or external events such as network activity.

#### MRML Scene

Is the workspace in Slicer. A Slicer scene is a MRML file that contains a list of elements (MRML Nodes) loaded into Slicer (volumes, models, fiducials). All data is stored in a MRML scene, which contains a list of MRML nodes. The in-memory state of the application can be serialized to a .mrml file at any time.

#### Pseudo-landmarks

Are generated automatically using the geometry of the 3D model without any reference to anatomically defined landmarks.

#### Segmentation

A data structure that stores the result of a segmentation (eg., ROI, mask, contour) and can be derived from one or more scalars volumes. The segmentation and the scalar volumes share a common spatial frame of reference, but do not need to be geometrically aligned or share the same voxel spacing. A segmentation can have various internal representations (binary labelmap, closed surface, etc.) which can be changed by the user. Some editing or analysis steps may require a particular representation. Segmentations can be exported as labelmaps or 3D models. Conversely, 3D models and label maps can be imported as segmentations. The segmentation data structure allows for overlap of segments; i.e., the same voxel can be assigned to multiple segments. This is different from label map representation, in which each voxel can only be assigned to a single class.

#### Semi-landmarks

Equidistant points on curves or patches which are drawn (curves) or calculated (patches) using an existing set of anatomical landmarks as boundaries or constraints.

#### Volume Rendering

Volume rendering (also known as volume ray casting) is a visualization technique for displaying image volumes as 3D objects directly - without requiring segmentation. This is accomplished by specifying color and opacity for each voxel, based on its image intensity. Several presets are available for this mapping, for displaying bones,soft tissues, air, fat, etc. on CT and MR images, which are typically calibrated to use Hounsfield units. Most microCT and other research imaging modalities are not calibrated, and when directly applied to such datasets, these presets will not perform well. However, users can fine tune the presets for their data by shifting the transfer function control points to the left or right.

#### Volume

3D array of voxels, such as CT, MRI, PET. It has different subtypes, such as:

1. **Labelmap volume**: each voxel value stores a discrete value, for example an index of a color in a color table; this can store segmentations (index value refers to segment number). This is the traditional output of a segmentation.
2. **Scalar volume**: each voxel stores a continuous value, such as intensity or density value (e.g., intensity values of a microCT)
3. **Vector volume:** each voxel contains a vector, can be used to store color (RGB) images, displacement fields, etc.

### SOM 4. SUPPLEMENTAL ANIMATION

This animation was created using the Animator module and data from https://github.com/muratmaga/Hominoid_Skulls/blob/master/Pongo_pgymaeus/Pongo_template_7_cleaned.nrrd?raw=true. Brain endocast was generated directly from the data using the SlicerMorph’s Segment Endocranium module applying the default settings. Lights module from the Sandbox extension was used to adjust the illumination and increase the depth perception.

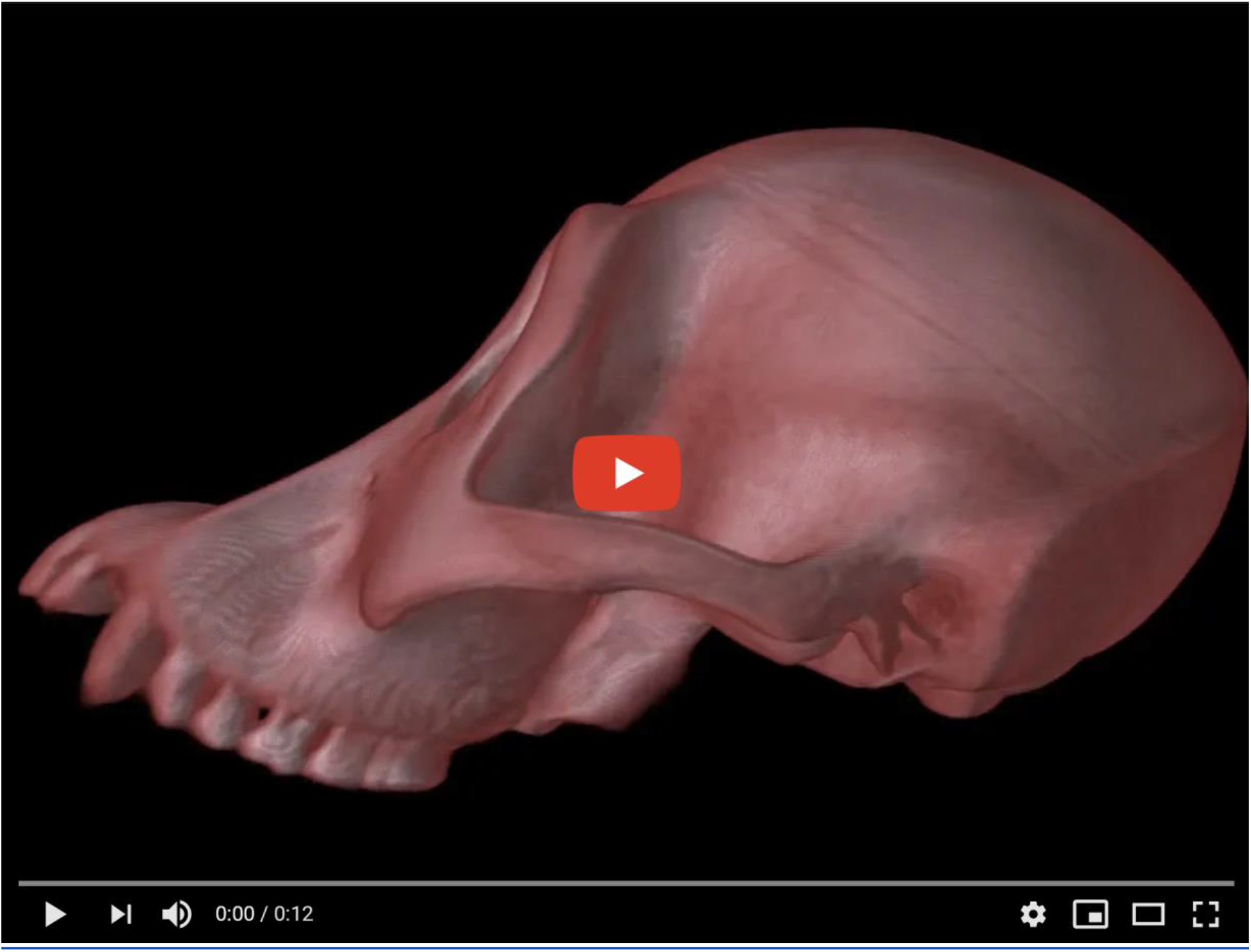

### SOM 5. IMPORTING 3D COORDINATES FROM SLICERMORPH INTO R

Traditionally, markups coordinates in Slicer were saved in FCSV format. FCSV is a derivative of comma-separated file format (csv) that contains three rows of header, which define the major Slicer version used to generate the data, coordinate system assumption and the column labels. 3D coordinates are stored in the fields labeled *x, y*, and *z*. Label associated with the landmark is stored in *label* field. SlicerMorph uses one landmark file per specimen paradigm, in which landmark filename will identify the specimen.

~~~
# Markups fiducial file version = 4.11
# CoordinateSystem = LPS
# columns = id,x,y,z,ow,ox,oy,oz,vis,sel,lock,label,desc,associatedNodeID
1, −35.628409103562,-7.275957614200383e-9,18.844778368826162,0,0,0,1,1,1,0,F-1,
2, −61.232510213366396,82.99982712156395,33.00776151257523,0,0,0,1,1,1,0,F-2,
3,17.265122462717727,52.50090683410033,-10.904087084733677,0,0,0,1,1,1,0,F-3,
~~~

Slicer community is in the process of switching to JSON, an industry standard lightweight generic data-interchange format, in lieu of FCSV. The benefit of JSON is the extensibility (additional tags can be added without breaking the format). While SlicerMorph supports FCSV for backward compatibility with existing data, we expect to switch to JSON as default markups output in upcoming releases. Below are two convenience functions that will import coordinate data from a markups file either in FCSV or JSON format. Examples of using the functions below to import GPA aligned or raw coordinates into R and conducting allometric regression directly on them is provided at: https://github.com/muratmaga/SlicerMorph_Rexamples/blob/main/geomorph_regression.R

#### R functions to import coordinates from SlicerMorph

~~~
# *Read in a .fcsv file from SlicerMorph*
read.markups.fcsv = function (file=NULL) {
 temp = read.csv(file=file, skip = 2, header = T)
 return (as.matrix (temp[, 2:4]))
}
# *Read in a mrk.json file from SlicerMorph*
read.markups.json = function(file=NULL) {
if (!require(jsonlite)) {
        print(“installing jsonlite”)
        install.packages(‘jsonlite’)
        library(jsonlite)
}
 dat = fromJSON(file, flatten=T)
 n=length(dat$markups$controlPoints[[1]]$position)
 temp = array(dim = c(n, 3))
 for (i in 1:n) temp[i,] = dat$markups$controlPoints[[1]]$position[[i]]
 return(temp)
}
~~~

## Notes

### Competing Interest Statement

The authors have declared no competing interest.

